# Triazole ureas covalently bind to strigolactone receptors and regulate signaling

**DOI:** 10.1101/362830

**Authors:** Hidemitsu Nakamura, Kei Hirabayashi, Takuya Miyakawa, Ko Kikuzato, Wenqian Hu, Yuqun Xu, Kai Jiang, Ikuo Takahashi, Naoshi Dohmae, Masaru Tanokura, Tadao Asami

## Abstract

Strigolactones (SLs), a class of plant hormones with multiple functions, mediate plant-plant and plant-microorganism communications in the rhizosphere. In this study, we developed potent strigolactone antagonists, which covalently bind to the strigolactone receptor D14, by preparing an array of triazole urea compounds. Using yeast two-hybrid assays and rice tillering assays, we identified a triazole urea compound KK094 as a potent inhibitor of strigolactone receptors. The LC-MS/MS analysis and X-ray crystallography concluded that KK094 was hydrolyzed by D14, and that a reaction product of this degradation covalently binds to the Ser residue of the catalytic triad of D14. We also identified KK052 and KK073, whose effects on D14–D53/D14–SLR1 complex formation were opposite due to a trifluoromethyl group on its benzene ring. These results demonstrate that triazole urea compounds are potentially powerful tools for agricultural application and may be useful for the elucidation of the complicated mechanism underlying SL-perception.

## MAIN TEXT

### Introduction

Strigolactones (SLs) are a class of plant hormones that control many aspects of plant growth and development. They promote the efficiency of arbuscular mycorrhizal symbiosis and the germination of parasitic plants of the Orobanchaceae family in the rhizosphere (*1*). Therefore, chemical regulators of SL functions might ultimately achieve widespread use in agricultural applications (*2*). To date, SL mimics have been developed to inhibit branching (*2, 3*) and trigger suicidal germination of parasitic weeds, and some compounds have succeeded in generating practical treatments that can induce suicidal germination of parasitic weeds in soil (*4*), and recently, several groups have reported the development of SL antagonists (*5–9*).

Important factors for SL perception and signaling have been identified using SL mutants (*10*), including those of D14, an α/β hydrolase (ABH) (*11, 12*), D3/MAX2 (*13, 14*), an F-box protein, and D53/SMXLs (*15–17*). It was recently postulated that SL is received by D14, which forms a complex with D3/MAX2 and D53/SMXLs (*16, 17*). This interaction stimulates the degradation of SL signaling inhibitors, including D53/SMXLs, which promotes the assembly of the TOPLESS corepressor-nucleosome complex (*18*), and enhances the expression of SL-inducible genes. Once SL is received by D14, it is hydrolyzed by D14 and a cleaved D-ring fragment is produced (*12, 19*). Recently, the D-ring-derived cleaved fragment was reported to form a covalent bond with a His residue in the catalytic center of D14 (*19, 20*). The formation of the D14-covalently linked intermediate molecule (CLIM) complex induces a dramatic change in the conformation of D14 and the formation of the D14–D3/MAX2 complex, which is essential for SL signaling (*21*). In addition, the SL hydrolysis of D14 is also required for the complex formation directly with D53/SMXLs (*16, 17*) and another target SLR1 (*19*), which is a negative regulator of gibberellin signaling. However, the mechanism of their SL-dependent interactions remains unclear.

Therefore, the blockade of the hydrolytic activity of D14 is a promising approach for the development of SL signaling inhibitors. D14 is an ABH, which belongs to a serine-hydrolase family. Serine-hydrolases universally possess a nucleophilic Ser-residue in their active sites, which is used for hydrolysis of their substrates. This nucleophilicity can be a target for covalent modification by reactive electrophiles. Indeed, wide-ranging types of electrophiles, including β-lactams/β-lactones, fluorophosphonates, and carbamates covalently modify the Ser-residue of target proteins (*22*). The tetrazole urea LY2183240-induced inhibition occurred via covalent carbamoylation of the serine nucleophile of FAAH (*23*). In addition, an isoxazolonyl urea and a 1,2,4-triazole urea were potent inhibitors of serine hydrolases (*24, 25*). These reports suggest that *N*-heterocyclic ureas, which can attack the nucleophilic Ser-residue in active sites of their targets, are a potent scaffold for serine hydrolase inhibitor design. Adibekian et al. synthesized a variety of 1,2,3-triazole ureas using click chemistry and determined that they selectively inhibit enzymes from diverse branches of the mammalian serine hydrolase family (*26*).

Here, we report that some 1,2,3-triazole ureas can inhibit the hydrolytic activity of the SL receptor D14 and stimulate rice tillering by blocking SL signaling. We also showed that D14 could degrade KK094, a potent 1,2,3-triazole urea D14-inhibitor. Using LC-MS/MS analysis and X-ray crystallography, we further confirmed that a reaction product of this degradation covalently binds to the Ser residue of the catalytic triad of D14. We also identified KK052 and KK073, whose effects on D14–D53 complex formation were opposite due to the presence or absence of trifluoromethyl group on the benzene ring. These compounds may have a potential as a useful tool to deepen our understanding of the molecular mechanism underlying SL-perception.

### Results

#### Synthesis and selection of candidate compounds for SL-receptor inhibitors by the yeast two-hybrid assay

According to the former report (*26*), we synthesized a series of 1,2,3-triazole ureas by mixing unsubstituted 1,2,3-triazole and five different carbamoyl chlorides with *N,N*-dimethyl-4-aminopyridine. This synthetic procedure yielded mixtures of N1- and N2-carbamoylated regioisomers. Most of these mixtures were separated by silica gel chromatography, and we obtained eight different 1,2,3-triazole urea congeners (Fig. 1A, **1**–**8**).

**Fig. 1.**
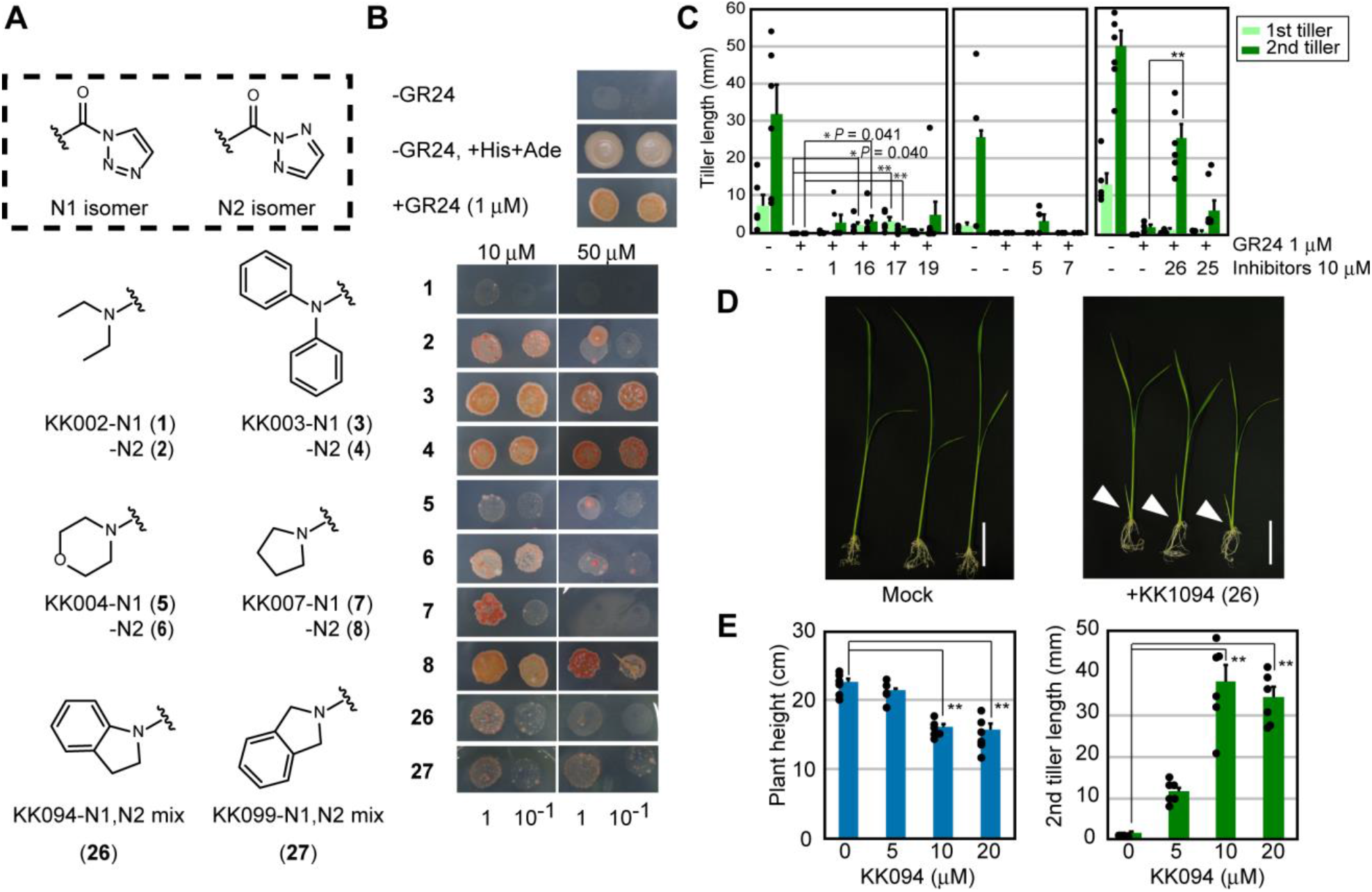
Inhibition of the SL-dependent D14-D53 interactions and rice tillering. (**A**) Structures of 1,2,3-triazole ureas. (**B**) Growth of AH109 yeast transformed with pGBK-D14 and pGAD-D53 were grown for 2 days in the liquid SD+His Ade media for 2 days at 30 °C and 5 μL of each yeast culture was spotted on SD-His Ade plates containing 1 μM GR24 with 10 μM or 50 μM concentrations of the 1,2,3-triazole ureas. The bottom numbers indicate dilutions of the yeast culture. (**C**) Inhibition of the effect of SL on rice tillering by 1,2,3-triazole ureas. Seven-day-old *d17-1* mutant rice seedlings were treated with experimental compounds and grown for a further 7 days, and the length of the first/second tillers were measured. (**D**) Stimulation of rice tillering by KK094. Eight-day-old wild-type rice (cv Nipponbare) seedlings were treated with/without 10 μM KK094 and grown for a further six days. Arrow heads indicate out growing tillers. Scale bars = 3 cm. (**E**) Concentration-dependent effects of KK094 on plant growth and rice tillering. Eight-dayold wild-type rice (cv Nipponbare) seedlings were treated with/without KK094 and grown for a further six days; and the length of first/second tillers were measured. Error bars indicate SE of six seedlings. Student’s *t*-test was used to determine the significance of differences (\**p*<0.05, \*\**p*<0.01).

To test whether these compounds are potent inhibitors of D14, we used the yeast two-hybrid (Y2H) assay to monitor SL-dependent D14–D53 interaction. Growth of AH109 yeast transformed with pGBK-D14 and pGAD-D53 in the SD+His Ade media were not inhibited by any KK compounds tested at the concentration of 50 μM showing that all KK compounds are not toxic for yeast growth at this concentration (data not shown). All tested compounds inhibited the D14–D53 interaction induced by GR24, a synthetic SL mimic, although the inhibitory effect of KK003 (**3**, **4**) was weaker than that of the other congeners (Fig. 1B).

To obtain highly specific and potent D14 inhibitors, we then diversified the structures of 1,2,3-triazole urea compounds. The 1,2,3-triazole urea scaffold consists of two major sites for diversification: the carbamoyl group and the 1,2,3-triazole leaving group. Adibekian et al. showed that 4-substitution of the 1,2,3-triazole group of 1,2,3-triazole ureas enhanced their potency and selectivity as serine hydrolase inhibitors (*26*). Therefore, we introduced various substituents onto the 4-position of the 1,2,3-triazole group of KK007, which clearly inhibited the D14–D53 and D14–SLR1 interactions induced by GR24 (fig. S1A, **9**–**15**). We tested these in the Y2H assay, but none of these compounds inhibited the D14–D53 and D14–SLR1 interactions induced by GR24 (fig. S1B), indicating that the unsubstituted 1,2,3-triazole is essential for the inhibition of D14 activity.

Next, we diversified our collection of D14 inhibitor candidates by converting carbamoyl groups. We synthesized 1,2,3-triazole urea compounds by modifying the morphorine structure of KK004 (fig S1C, **16**–**25**), and the pyrrolidine structure of KK007 (Fig. 1A, **26**–**27**). All of these were tested in the Y2H assay (Fig. 1D and fig. S1D). As a result, we selected KK002-N1, KK004-N1, KK007-N1, KK052, KK053, KK055, KK075, and KK094 for further assays.

#### Inhibitory effects of candidate compounds on the suppression of rice tillering by GR24

To confirm whether the selected compounds are good SL-receptor inhibitor candidates, we used *in planta* assays. In hydroponically grown rice seedlings, the first and second tiller buds of SL-deficient mutants, such as *d10* and *d17*, grow out, whereas those of wild-type plants remain dormant (*27*). Treatment of these mutants with SLs restores the dormant phenotype of the first and second tiller buds. Therefore, we performed a rice tillering assay to test whether 1,2,3-triazole ureas attenuate the tiller-inhibiting effects of SLs. When 10 μM of 1,2,3-triazole ureas and 0.1 μM of GR24 were applied together to hydroponically grown *d17-1* rice, several 1,2,3-triazole ureas restored the growth of the first and second tiller bud that had been suppressed by GR24 (Fig. 1C). KK052 (**16**), KK053 (**17**), and KK094 (**26**) significantly restored the tiller bud outgrowth of rice (Fig. 1C). Among them, the effect of KK094 treatment was prominent. KK094 did not inhibit the growth of rice, while KK052 did (fig. S2). We then substituted the 1,2,3-triazole group of KK094 to 1,3-imidazole group to yield KK122 and tested it using the Y2H assay and the rice tillering assay. KK122 did not inhibit the GR24-induced D14–D53 and D14–SLR1 interactions, or the tillering inhibition produced by GR24 (fig. S3). These data indicate that the 1,2,3-triazole group is indispensable for the inhibition of D14 function. Next, we generated various KK094 derivatives by modifying the indolinyl structure of KK094 (fig. S4, **29**–**41**); however, none of these was a stronger inhibitor than KK094. Thus, we selected KK094 as the most potent SL-receptor inhibitor candidate, and KK122 as a negative control compound of KK094.

We then treated hydroponically grown seedlings of wild-type rice (Nipponbare) with 10 µM KK094 and found that it significantly promoted the outgrowth of the first and second tiller buds (Fig. 1D), and this outgrowth promotion was concentration-dependent (Fig. 1E). The KK094 treatment decreased plant height. A semi-dwarf phenotype is a characteristic of SL-deficient mutants. These features resulting from KK094 treatment support the hypothesis that this compound acts as an SL-signaling inhibitor in rice. Furthermore, we investigated whether KK094 can inhibit the germination of parasitic plants and found that it could inhibit GR24-induced seed germination of *Striga hermonthica* (fig. S5A).

#### KK094 inhibits SL-hydrolysis

Because KK094 was originally designed to inhibit the SL-hydrolytic activity of D14 by direct binding to its catalytic center, we tested whether KK094 inhibits D14-induced SL-hydrolysis. When 0.2 μM of GR24 was incubated with the purified recombinant D14 for 30 min, and GR24 and the reaction product, ABC-OH, were extracted and detected by LC-MS, a decrease in the amount of GR24, and an increase in the amount of ABC-OH were observed (fig. S6A–C). We next conducted the SL-hydrolysis assay at various concentrations of KK094. As a result, KK094 efficiently inhibited the decrease in the amount of GR24, and the increase in the amount of ABC-OH produced by D14, and these inhibitory effects were concentration-dependent (Fig. 2A). KK094 itself was also hydrolyzed by D14 (fig. S6D), indicating that the hydrolyzed product may bind to the catalytic pocket of D14 and impede entry of SL into the pocket.

**Fig. 2.**
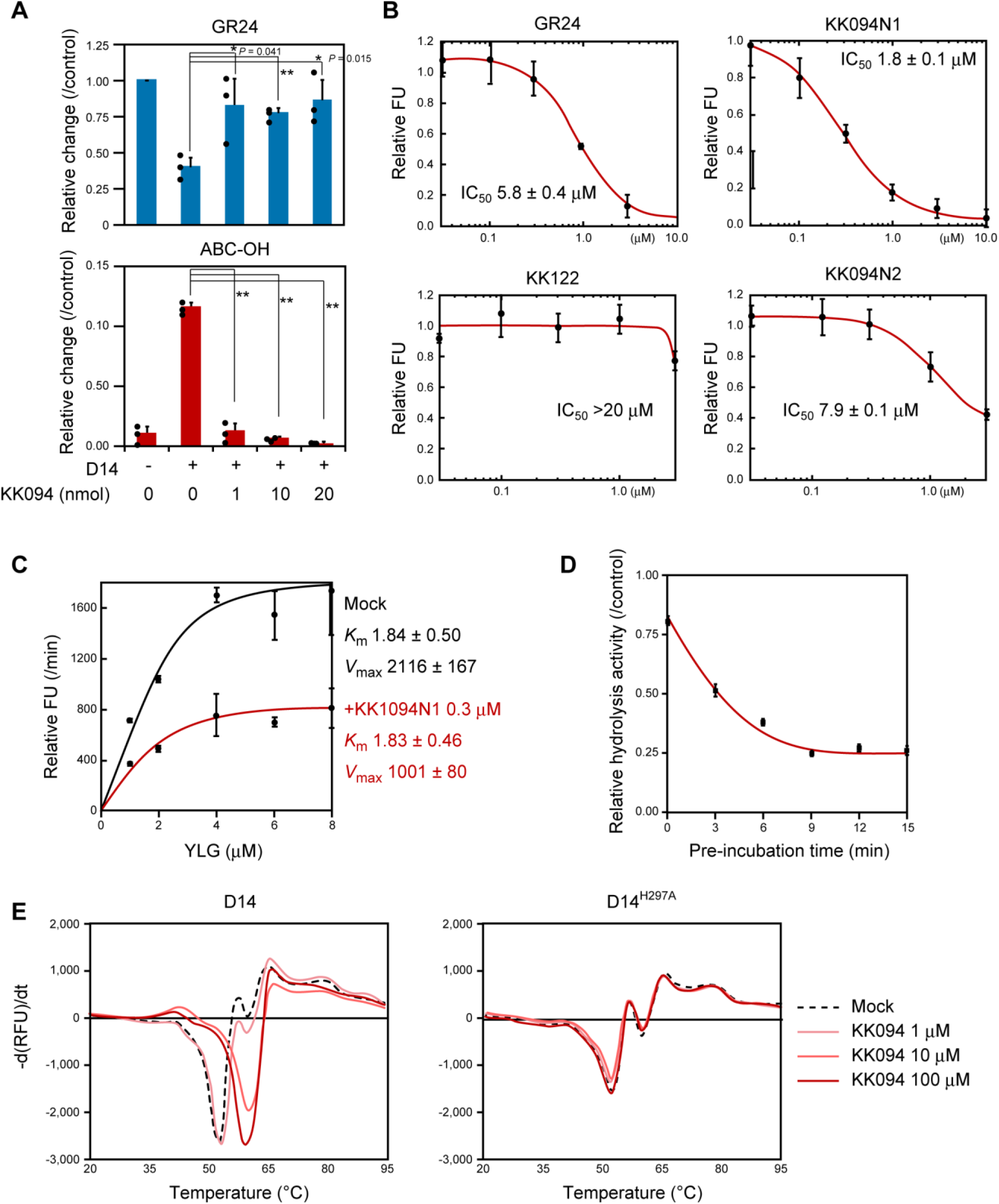
Inhibition of D14 by KK094. (**A**) KK094 concentration-dependently inhibits hydrolysis of GR24 by D14. GR24 and KK094 were pre-incubated at 30° C for 10 min and incubated further at 30° C for 20 min after addition of 0.33 μM of D14. Then the quantities of GR24 or ABC-OH were monitored by LC-MS. The amount of GR24 incubated without D14 was set to 1. Error bars indicate SE of three sample replicates. Student’s *t*-test was used to determine the significance of differences (\**p*<0.05, \*\**p*<0.01). (**B**) Competitive inhibition of D14-mediated Yoshimulactone G (YLG) hydrolysis. A concentration of 0.3 μM of YLG was reacted with 0.33 μM of D14 and the fluorescent intensity was measured at an excitation wavelength of 485 nm and detected at 535 nm. The fluorescent values relative to the hydrolysis without inhibitor was shown. FU = fluorescence unit. Error bars indicate SE of three sample replicates. IC_50_ values were calculated using the website http://www.ic50.tk/index.html. (**C**) KK094 alters the *V*_max_ value, not the *K*_m_ value, of YLG hydrolyzation by D14. Error bars indicate SE of three sample replicates. 0.3 μM of KK094-N1 reduced the *V*_max_ value by half but did not change the *K*_m_ value. (**D**) Preincubation with KK094-N1 reduced hydrolysis activity of D14. Pre-incubation of 0.33 μM of D14 with 0.3 μM of KK094-N1 for the indicated time and incubated for another 10 min after the addition of 0.3 μM of YLG, and the fluorescent intensity was measured. Error bars indicate SE of three sample replicates. (**E**) Melting temperature curves for D14 and the D14^H297A^ mutant protein at varying concentrations of KK094 as monitored by differential scanning fluorometry. Each line represents the average protein melt curve for three sample replicates measured in parallel.

Yoshimulactone Green (YLG) is a fluorogenic SL-agonist which suppresses the more axillary branching phenotype of the *Arabidopsis max4* mutant and stimulates the seed germination of *Striga* (*28*). YLG can be efficiently hydrolyzed by AtD14 and *Striga hermontica* SL receptors (ShHTLs). YLG-hydrolysis by SL receptors generates the fluorescent product. First, we observed that recombinant D14 protein hydrolyzed YLG to yield green fluorescence, which over time showed a Michaelis constant (*K*_m_^YLG^) value at 1.84 μM (Fig. 2C). This reaction was competitively inhibited by GR24, and the median inhibitory concentration (IC_50_) was 5.8 μM (Fig. 2B). Then we investigated whether KK094 competes with YLG in its hydrolysis by D14. KK094 is a mixture of regioisomers, KK094-N1 and KK094-N2, which we separated using silica gel chromatography. Both the N1 and N2 isomers of KK094 inhibited YLG hydrolysis by D14, with IC_50_ values of 1.8 and 7.9 μM, respectively (Fig. 2B). KK122 did not inhibit YLG hydrolysis (Fig. 2B), indicating that the 1,2,3-triazole moiety is essential for the inhibition of YLG hydrolysis and that the inhibition of YLG hydrolysis is required for the inhibition of SL-signaling. The *K*_m_^YLG^ value was unchanged in the presence of KK094-N1 at a concentration of 0.3 μM. However, KK094-N1 treatment at a concentration of 0.3 μM decreased the *V*_max_^YLG^ to nearly half the level observed following sham treatment (Fig. 2C), and the increase of KK094-N1-pretreatment time potentiated the inhibitory effect (Fig. 2D). These data strongly support the hypothesis that KK094 is a covalent inhibitor of D14.

We also assessed the inhibitory effect of KK094 on SL-hydrolysis activity of *Striga* SL-receptors using YLG. *S. hermonthica* receives SL by the ShHTL protein, of which there are 11 subtypes, upon SL-induced seed germination. Among them, the ShHTL7 subtype is reported to be the most sensitive SL receptor (*29*) mediating SL-induced seed germination, therefore, we used the ShHTL7 protein in the YLG assay. The IC_50_ of KK094 was more than 20 μM, while that of GR24 was 0.15 μM (fig. S5B).

Next we monitored the interaction between KK094 and D14 using differential scanning fluorimetry (DSF). DSF revealed a shift in the D14 melting temperature in the presence of KK094, indicating that KK094 interacts with D14 and changes its stability; however, with KK094 treatment, this shift was not observed in D14^H297A^ in which the catalytic residue is mutated (*19*) (Fig. 2E). The 8 °C increase in D14 melting temperature suggests that the stabilization of D14 and the potent inhibitory effect of KK094 are correlated.

#### KK094 inhibits SL-induced formation of the D14–D53 complex *in vitro*

We performed *in-vitro* pull-down assays (Fig. 3A) using a two-step treatment of D14 with test compounds; D14 was treated with the first compound, and then washed and incubated with D53 in the presence of the second compound. The single treatment with KK094 did not induce the D14–D53 interaction when it was added in either step. Meanwhile, the single treatment with GR24 did not induce the interaction when it was added in the first step, but it did only when added in the second step, indicating that a hydrolyzed product of GR24 could be washed away from the catalytic pocket. However, when D14 was treated with KK094 (first step) and GR24 (second step), the interaction was not detected, indicating that a hydrolyzed product of KK094 stayed in the pocket and inhibited GR24 hydrolysis, and the hydrolyzed product was not washed away in the washing step. These results strongly support a mechanism of covalent inhibition by KK094. A concurrent treatment with KK094 and GR24 weakened the interaction between D14 and D53 as compared with GR24 alone.

**Fig. 3.**
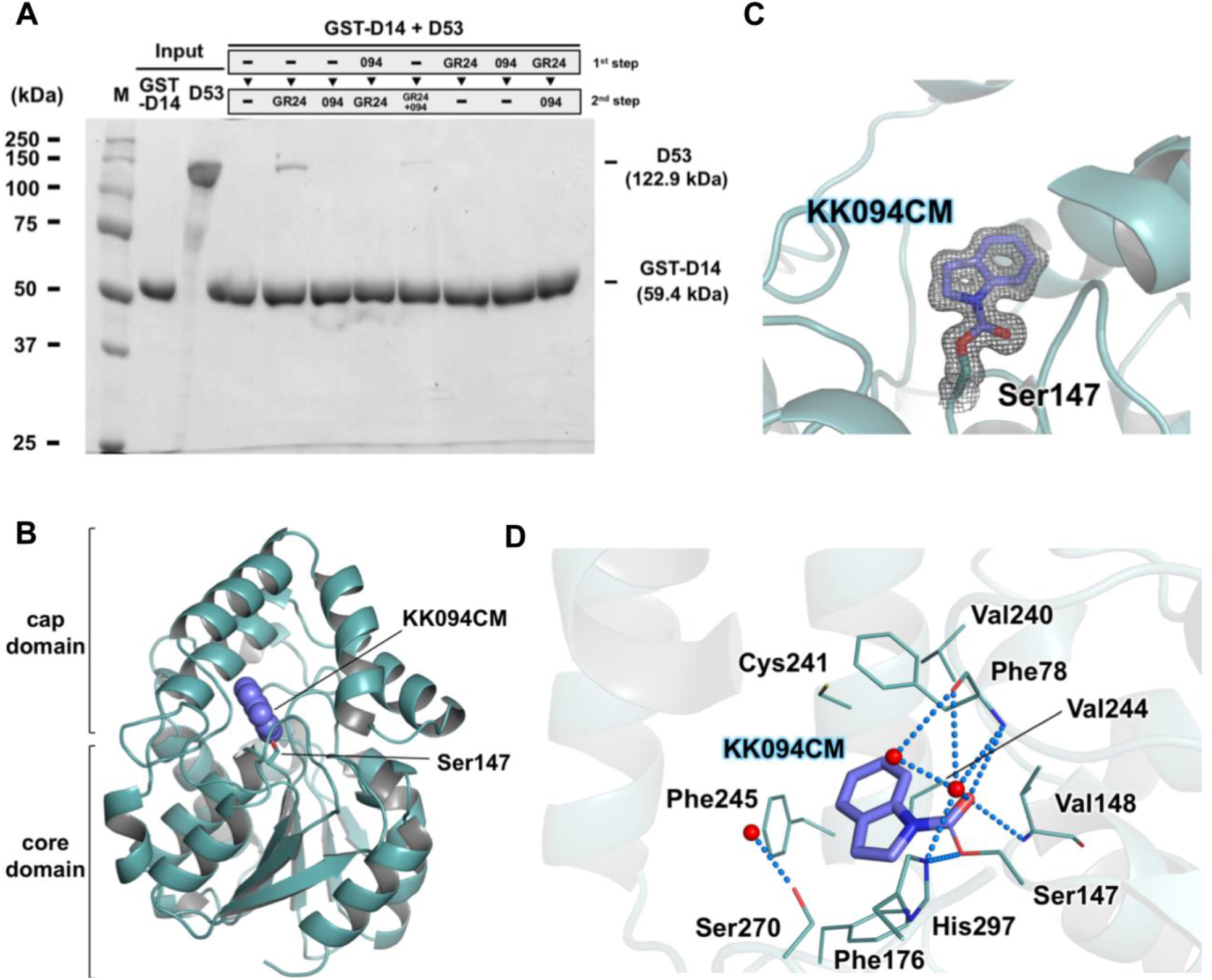
*in-vitro* pull-down assays of GST–D14 and D53 proteins, and the crystal structure of D14 in complex with KK094CM. (**A**) The *in-vitro* pull-down assays were performed using a two-step treatment with GR24/KK094. The interaction of D14 with D53 was detected by SDS-PAGE and Coomassie Brilliant Blue (CBB) staining. (**B**) Overall structure of KK094CM-bound D14. The covalently bound KK094CM is depicted in purple with van der Waals surfaces, and the catalytic S147 residue is shown as a stick model. (**C**) The electron density map of the covalently bound KK094CM to S147. KK094CM is shown as a stick model (purple), along with the 2*F*_o_-*F*_c_ map contoured at 1.0σ (gray). (**D**) Binding site of KK094CM in the catalytic pocket of D14. The residues involved in the interaction with KK094CM (< 4.5 Å) are represented by a line model. Water molecules are shown as red balls, and the blue broken line indicates a hydrogen bond.

#### Crystal structure of the D14–KK094 complex

To determine the binding mode of KK094 in the catalytic pocket of D14, we solved the crystal structure of the D14–KK094 complex at 1.45 Å resolution (table S1, Fig. 3B). The asymmetric unit contains two molecules of D14 with almost identical structures; the root mean square deviation (r.m.s.d.) was 0.37 Å for the main chain Cα atoms. As reported previously, D14 consisted of a core domain, also known as α/β hydrolase domain (*30*), and a cap domain composed of four helices forming two antiparallel V shapes (Fig. 3B). A catalytic pocket was formed between the two domains, and the catalytic residue S147 is located at the bottom of the pocket and is aligned with H297 and D268 to form the catalytic triad. The electron density map of the D14–KK094 complex clearly showed the existence of a covalently bound KK094-derived carbamoyl moiety (KK094CM), which is assumed to be a hydrolyzed product of KK094, to the hydroxyl group of S147 (Fig. 3C). In the complex structure of D14–KK094CM, the covalently bound KK094CM was embedded completely in the cavity and was surrounded by V148, V240, C241, V244, S270 and H297, and several aromatic residues, such as F78, F176 and F245 (Fig. 3D). These residues made favorable hydrophobic and/or van der Waals interactions with KK094CM. In addition, the carbonyl group of KK094CM formed a hydrogen bond network with F78, V148, and H297 including two water molecules (Fig. 3D). In this complex structure, KK094CM is deduced to deprive canonical substrates of opportunities to invade and be hydrolyzed by occupying the active site in the catalytic pocket of D14. The D14–KK094CM complex and the apo-D14 (PDB ID 3VXK) (*19*) have almost identical structures (r.m.s.d. 0.16 Å), but some minor structural differences occurred in the catalytic pocket (fig. S7A). The side chain of S147 flipped toward KK094CM due to the covalent modification. In addition, upon KK094CM binding, the side chain of F245 moved 1 Å away from the KK094CM, creating a space to accommodate KK094 binding (fig. S7B). This change is consistent with previous structural studies regarding the flexibility of the Phe residue located in this conserved site (*19, 31–33*).

#### Hydrolyzed product of KK094 covalently binds to D14

We further confirmed whether KK094 covalently binds to D14 using Matrix Assisted Laser Desorption/Ionization-Time of Flight (MALDI-TOF) mass spectrometry and Liquid Chromatography-Tandem Mass Spectrometry (LC-MS/MS) analysis. D14 was incubated with or without KK094 in 1 mL of PBS at 25 °C for 35 min, and 100 μL of the reaction solution was analyzed using MALDI-TOF mass spectrometry (Fig. 4A). The molecular weight of D14 incubated without KK094 was determined to be 29283.3 (-KK094; Fig. 4A), which was almost the same as that deduced from the amino acid sequence of the purified D14 protein (molecular weight: 29285.8). The molecular weight of D14 incubated with KK094 was determined to be 29429.7 (+KK094; Fig. 4A). The 146.4 Da difference was thought to represent the covalently bound KK094CM (Fig. 4A), which adds 145 Da.

**Fig. 4.**
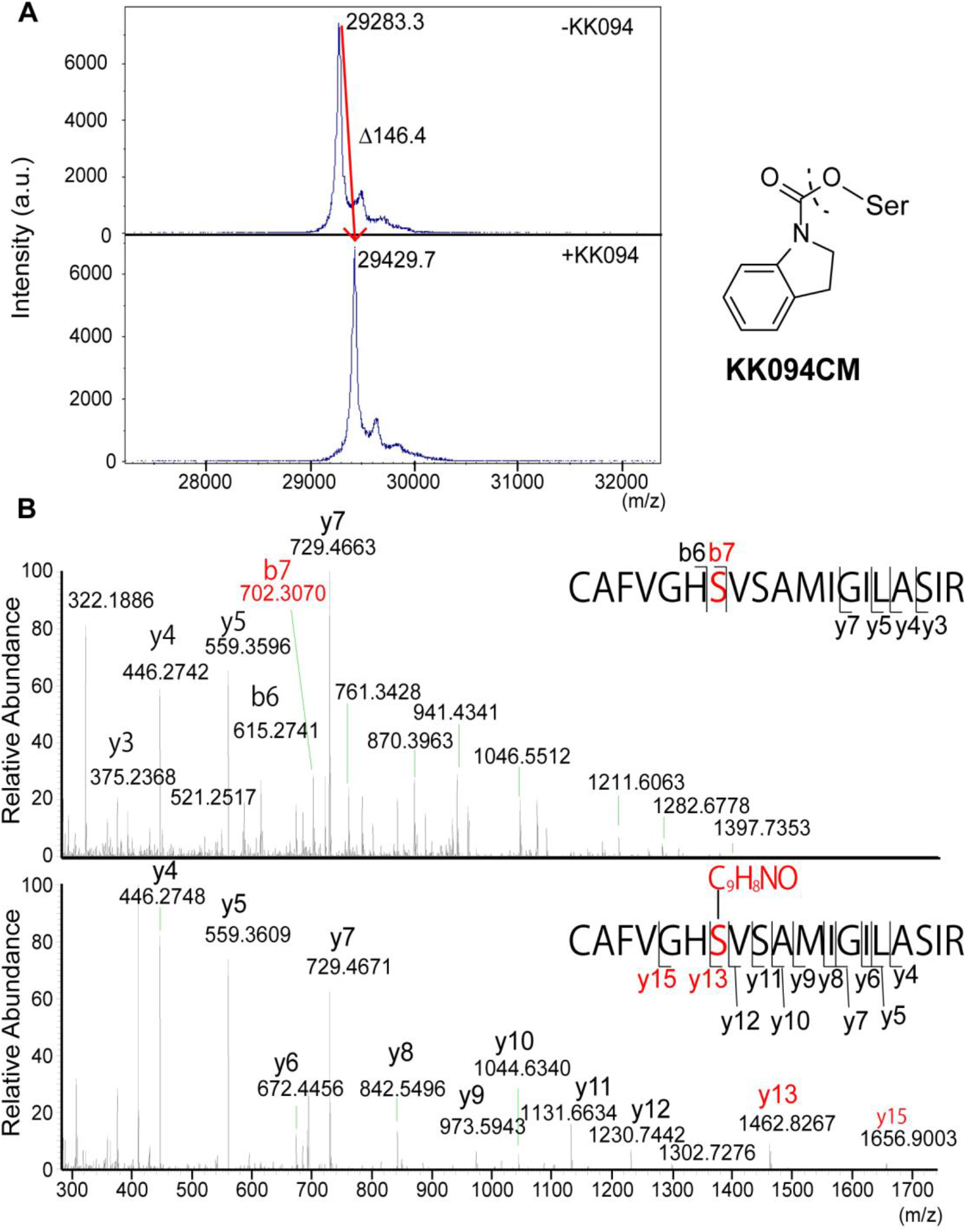
Covalent binding of KK094CM to S147. (**A**) D14 was incubated without (upper panel) or with (lower panel) GR24 and analyzed by MALDI-TOF MS. (**B**) The reaction mixture of D14 and GR24 without or with KK094 (-KK094/+KK094, respectively) was dialyzed against a 50 mM NH_4_HCO_3_ solution. The dialyzed sample was digested with trypsin and analyzed using LC-MS/MS. MS/MS spectra of peptides corresponding to peptides 141–159 (^141^CAFVGHSVSAMIGILASIR^159^; *m*/*z* 1932.446 for -KK094 (upper), *m*/*z* 2077.454 for +KK094 (lower)) are shown. Peaks of the ions y13 and y15 indicate that the Ser-residue (red letter) is covalently modified by KK094CM.

The remaining reaction mixtures were then dialyzed and digested with trypsin, and generated peptides were analyzed using MALDI-TOF mass spectrometry (fig. S8A). The amino acid sequences of the digested peptides were deduced based on their molecular masses (table S2). The fragment corresponding to the peptide 141–159 (^141^CAFVGHSVSAMIGILASIR^159^; *m*/*z* 1932.446) was not detected when D14 was incubated with KK094 (fig. S8B). On the contrary, the peptide fragment with molecular mass of 2077.454 was detected when D14 was incubated with KK094, although it was not detected when D14 was incubated without KK094. This fragment corresponds to the peptide 141–159 with covalently bound KK094CM (fig. S8B).

To confirm the carbamoylation site in peptide 141–159, we conducted LC-MS/MS analysis (Fig. 4B). When trypsin-digested fragments of D14 incubated with KK094 were analyzed, the addition of 145 Da was observed in the y13 and y15 ions but not in the y4–y12 ions (Fig. 4B; table S3), demonstrating that carbamoylation occurred at S147.

#### Some KK052-derived 1,2,3-triazole ureas show agonist activity in the formation of the D14–D53 complex

As described above, KK052 acted as an SL-antagonist in Y2H and rice tillering assays (fig. S1C and D), but its inhibition seemed weaker than that of KK094 (Fig. 1A). Two KK052-derivatives, KK053 and KK055, showed antagonism both in the Y2H and rice tillering assays (fig. S1C, D and Fig. 1C). Unexpectedly, some KK052-derivatives, KK054, KK067, KK070, and KK073 showed agonism in the formation of the D14–D53/D14–SLR1 complex in the Y2H assay (fig. S9 and Fig. 5A). Interestingly, these agonists had polar groups. Among them, the agonistic effect of KK073 was prominent (Fig. 5A). Thus, we further analyzed the activity of KK073.

**Fig. 5.**
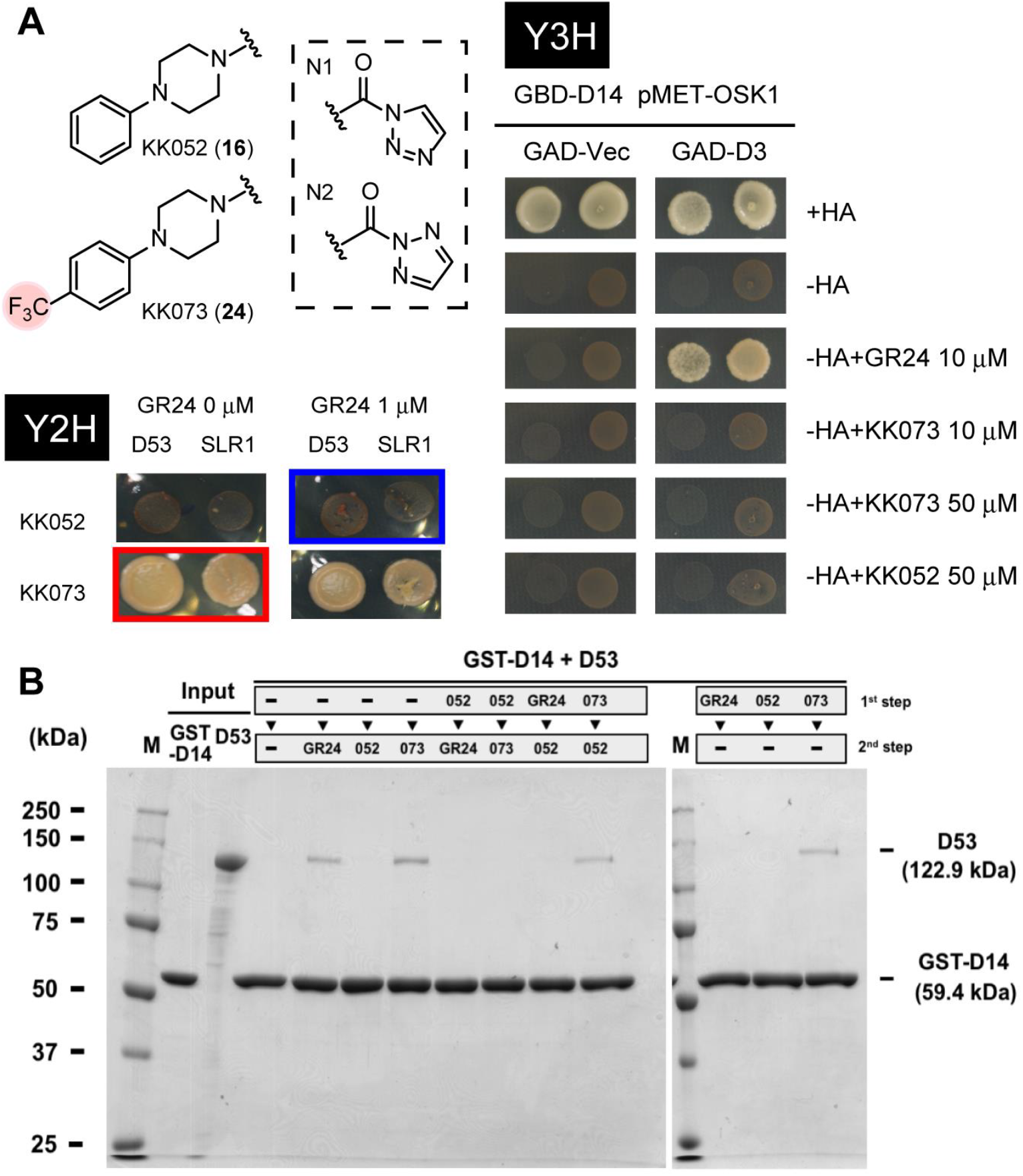
Agonistic effects of KK073 on D14–D53/D14–D3 interactions. (**A**) Structures of KK052 and KK073 are shown on the right. Y2H: Growth of AH109/pGBK-D14–pGAD-D53 or AH109/pGBK-D14–pGAD-SLR1 on SD-His Ade plates containing experimental compounds for 4 days at 30°C. KK073 showed an agonistic effect (in red squares) and KK052 showed antagonism (in blue squares). Y3H: Growth of AH109/ pBridge-BD:D14-M:OSK1–pGAD-D3 or AH109/ pBridge-BD:D14-M:OSK1–pGADT7 on SD-His Ade Met plates containing experimental compounds for 4 days at 30°C. (**B**) *In-vitro* pull-down assays of GST–D14 and D53 using a two-step treatment with GR24/KK052/KK073. The interaction of D14 with D53 was detected by SDS-PAGE and CBB staining.

#### KK052 inhibits the SL-induced formation of the D14–D53 complex and KK073-induced D14–D53 complex formation *in vitro*

We further conducted *in vitro* pull-down assays using GR24, KK052, and KK073 (Fig. 5B). Like the pull-down assay using KK094, we also used a two-step treatment of D14 with this group of compounds. A single treatment with KK052 did not induce the D14–D53 interaction when it was added in either step. As observed for the KK094 treatment (Fig. 3A), when D14 was treated with KK052 (first step) and then GR24 (second step), the D14–D53 interaction was not detected, indicating that a hydrolyzed product of KK052 (KK052CM) stayed in the pocket and inhibited the action of GR24. Meanwhile, the interaction between D14 and D53 was induced when KK073 was added in either step (Fig. 5B). When a combination of KK073 and KK052 was added, the interaction was induced only when KK073 was added in the first step. These results indicate that a hydrolyzed product of KK073, KK073CM, is not washed away from the catalytic pocket and promotes the formation of the D14–D53 complex by a mode of agonistic action different from GR24.

#### The hydrolyzed products of KK052 and KK073 covalently bind to D14

To determine whether KK052 and KK073 also covalently bind to D14, we solved the crystal structures of the D14–KK052 and the D14–KK073 complex at 1.49 Å and 1.53 Å resolution, respectively (table S1 and Fig. 6A). The electron density map of the D14–KK052 and D14–KK073 complex clearly showed the existence of a covalently bound KK052CM or KK073CM, respectively, to the hydroxyl group of S147 (Fig. 6B). In these complex structures, the covalently bound KK052CM was embedded completely within the cavity, whereas the trifluoromethyl group of KK073CM partially protruded out of the pocket. Both binding sites consisted of V148, I191, V194, C241, V244, S270 and H297, and several aromatic residues, such as F78, F176, F186, W205, Y209, and F245 (Fig. 6C). These residues made favorable hydrophobic and/or van der Waals interactions with KK052CM/KK073CM. In addition, the carbonyl group of KK052CM/KK073CM formed a strong hydrogen bond network with F78, V148, Y209, S270 and H297 mediated by three water molecules (Fig. 6C). Like KK094CM, KK052CM/KK073CM appears to deprive canonical substrates of opportunities to invade and react by occupying the active site. Interestingly, when the structure of the D14–KK073CM complex was compared to that of the D14–D-OH complex (PDB ID 3WIO) (*19*), the trifluoromethyl group of KK073 resided at nearly the same site as the D-OH in the D14–D-OH complex (Fig. 6D). Both the trifluoromethyl group of KK073 and D-OH were located at the aperture of the binding pocket of D14 and surrounded by aromatic residues, such as F186, W205, Y209, and F245. In addition, the trifluoromethyl group of KK073 protruded from the binding pocket and was directly exposed to the solvent; fluorine atoms faced on the surface of D14, generating a polar region in the overall hydrophobic surface of D14 (Fig. 6E). To confirm this hypothesis, we tested KK182, in which a methylene group was inserted between the benzene ring and the piperazine ring, and the distance between the trifluoromethyl group and the carbonyl group is elongated. When KK182 was tested in the Y2H assay, KK182 antagonized the formation of D14–D53 (fig. S9B). These data support the hypothesis that the position of the trifluoromethyl group is critical for the agonistic effect on the formation of the D14–D53/D14–SLR1 complex.

**Fig. 6.**
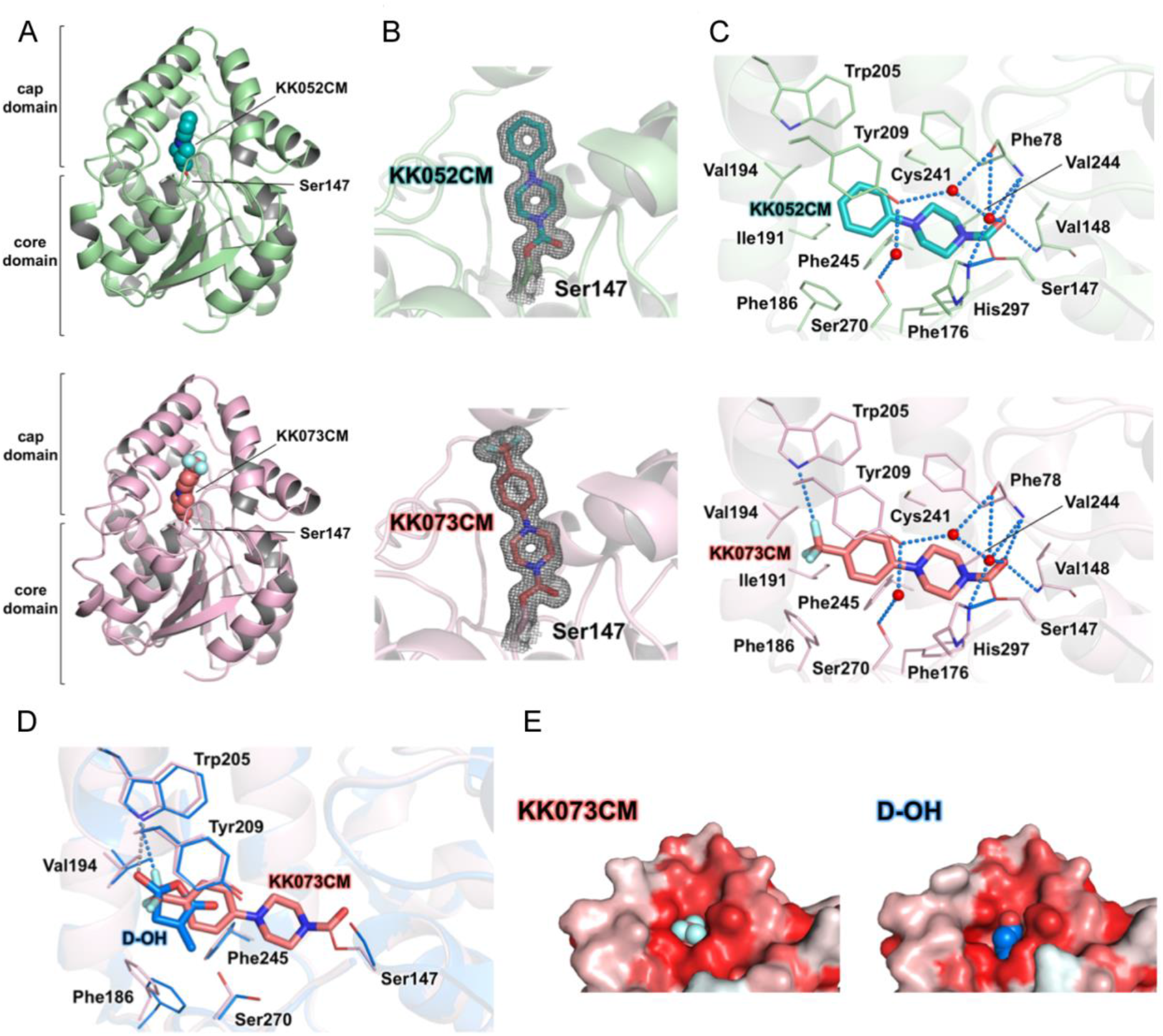
KK052CM and KK073CM covalently binds to D14. (**A–C**) Crystal structures of D14 in complex with KK052CM or KK073CM. Overall structures (**A**), electron density maps (**B**) and binding sites (**C**) of KK052CM (green) and KK073CM (pink). The orientation and structural representation are the same as in Fig. 3**A–D**. (**D**) Comparison of the ligand binding sites of KK073CM-bound D14 (pink) and D-OH-bound D14 (PDB ID 3WIO) (blue). The residues involved in the interaction with the trifluoromethyl group of KK073 and D-OH are represented by line models, and the broken lines indicate hydrogen bonds. The orientation is the same as in (**C**). The r.m.s.d. for the Cα atoms was 0.38 Å. (**E**) The hydrophobicity of the surface residues in KK073CM-bound D14 and D-OH-bound D14 (PDB ID 3WIO). Hydrophobic residues are shown in red, whereas hydrophilic sites are shown in white. KK073CM and D-OH are depicted in van der Waals surfaces. The trifluoromethyl group of KK073 protruded out of the pocket and generated a polar patch in the hydrophobic surface of D14.

#### Both KK052 and KK073 antagonized the inhibition of rice tillering produced by SL

From the above results, we assumed that KK052 acts as an SL-antagonist, and KK073 acts as an SL-agonist *in planta*. To confirm this assumption, we tested KK052 and KK073 in the rice tillering assay. Unexpectedly, both KK052 and KK073 showed inhibitory effects on the tillering inhibition by GR24, and KK073 did not act as an SL-agonist (fig. S10A). Generally, SL-agonists decrease the D14 melting temperature (*12, 20*), which is required for the conformational change in the interaction with D3/MAX2. However, KK073 as well as KK094 (Fig. 2E), increased the melting temperature (fig. S10B), indicating that KK073 interacts with D14 and changes the stability of D14, and this shift was not observed in D14^H297A^. Consistent with this, we found that KK073 could not induce the interaction of D14 with the D3–OSK1 complex in the yeast three-hybrid assay (Fig. 5A).

### Discussion

The irreversible mode of inhibition of covalent inhibitors has potential benefits, such as high potency extended duration of action as compared with that of the reversible mode of inhibition of noncovalent inhibitors. In this study, we reported the development of 1,2,3-triazole ureas with inhibitory effects on SL activity. Among them, KK094 was the most potent SL-inhibitor. KK094 restored the growth of the first and second tiller bud of hydroponically grown *d17-1* rice that has been suppressed by GR24 (Fig. 1C). KK094 also showed an inhibitory effect on the regulation of tiller bud growth by SLs and the treatment of wild-type rice with KK094 stimulated the growth of tiller buds (Fig. 1D, E). KK094 efficiently inhibited the hydrolysis of GR24 by D14, and this inhibition did not change the *K*_m_ value, but markedly lowered the *V*_max_ value of the hydrolysis reaction (Fig. 2). Pull down analyses showed that KK094 prevents the formation of D14–D53 complex *in vitro* (Fig. 3). KK094 could be hydrolyzed by D14 (fig. S6D), and the hydrolyzed product (KK094CM) covalently binds to the hydroxyl group of a Ser residue in the catalytic triad of D14, as indicated by the crystal structure analysis (Fig. 3B-D) and the LC-MS/MS analysis (Fig. 4). Based on these data, we conclude that KK094 is a potent covalent inhibitor for the strigolactone receptor.

Recently, Xiang et al. reported that β-lactones covalently bind to Arabidopsis D14 and inhibit the activity of AtD14 as an SL receptor (*7*). Here, we used 1,2,3-triazole ureas because they are easily synthesized using a simple scheme. This is a great advantage of 1,2,3-triazole ureas, and we synthesized a wide variety of 1,2,3-triazole urea compounds and tested them both using *in planta* and *in-vitro* assays and found some compounds with a spectrum of activities. Among them, we are interested in the compounds which seemed to act as agonists for the formation of D14–D53/D14–SLR1 complex in Y2H assays.

In our Y2H assays, KK052 inhibited SL-induced formation of the D14–D53 complex, while KK073 induced formation of the D14–D53 complex. This agonistic effect of KK073 was also observed in the pull-down assay. The difference between KK052 and KK073 is the existence of a trifluoromethyl group on the benzene ring. The opposite activity of KK052 and KK073 resulted from the slight difference which provides clues to a deeper insight into the mechanism of the formation of the D14–D53 complex.

In the pull-down assay, KK094 inhibited SL-induced formation of the D14–D53 complex by forming a covalent bond with the catalytic S147 residue. This result indicates that the complex formation requires SL binding into the catalytic pocket followed by its hydrolysis. The previous report showed that the hydrolyzed product of SL covalently bound to the catalytic site of D14 (*20,21*). They showed that this binding evoked dramatic changes in the overall structure of D14 and induced D14–MAX2 complex formation (*21*). In contrast, SL-induced formation of the D14–D53 complex could not be detected following a washing step of D14 in complex with SL before the incubation with D53, suggesting that the hydrolyzed product of SL binds non-covalently to D14 and induces the formation of the D14–D53 complex. In our previous report, we observed the non-covalently bound D-OH in the entrance of the ligand binding pocket of D14. It is possible that this D-OH induces the formation of the D14–D53 complex in a way different from that in the formation of D14–MAX2 complex.

Unlike GR24, KK073 could induce the formation of the D14–D53 complex even when it was added before the washing step in the pull-down assay. Furthermore, we observed covalent binding of KK073CM to the S147 residue of D14 using X-ray crystallography. Interestingly, the trifluoromethyl group of KK073CM in the D14–KK073CM crystal resided at almost the same site as the D-OH in the D14–D-OH crystal (Fig. 8). Both the trifluoromethyl group of KK073 and D-OH were located at the aperture of the D14 binding pocket. No major change of the overall structure of the complex was observed in both the D14–D-OH (*19*) and D14–KK073CM complexes (fig. S7).

KK052 could not induce the D14–D53 interaction, while KK073 could function as an agonist for the formation of the D14–D53 complex. The position of KK073CM in the binding pocket of D14 was almost the same as that of KK052CM, and the only difference is the trifluoromethyl group on the benzene ring of KK073CM. These data strongly indicate that the formation of the D14–D53 complex requires an additional polar region (e.g., D-OH, trifluoromethyl group) at the entrance of the binding pocket of D14. In addition, there are no structural differences in D14 between the KK073CM- and KK052CM-bound forms, suggesting that the formation of the D14–D53 complex does not require the structural changes that are observed in the D14–D3/MAX2 complex. Indeed, we found that KK1073 could not induce the formation of the D14–D3/MAX2 complex in the yeast three-hybrid assay (Fig. 5C). Furthermore, KK073 increased the D14 melting temperature in the DSF assay (fig. S10B) as well as KK094, while SL-agonists generally decrease the D14 melting temperature (*12,20*). In general, the decrease of the melting temperature reflects the destabilization of protein that correlates with the conformational change, which is consistent with the dynamic structural changes of D14 in the complex with D3/MAX2 with SL-agonists. In the D14–D3/MAX2 complex, the reposition of the catalytic triad accompanying the formation of CLIM may induce the destabilization of D14 (*21*). On the other hand, covalently bound KK094CM/KK073CM interacts with many residues composing the pocket and is likely to decrease their flexibility, resulting in the stabilization of D14. These data also support a model of the static formation of the D14–D53 complex.

These speculations raise the question of how D14 interacts with both D53 and D3/MAX2 by different means. We do not have any direct evidence to answer this question. Investigation of the structure of the complex containing D14, D53 and D3/MAX2 might provide insights regarding this question.

Although KK094 strongly inhibited the SL activity in the suppression of the outgrowth of branching buds, the inhibitory effect of KK094 on the SL-induced germination of *Striga* seeds was relatively weak (fig S5). A greater KK094/GR24 ratio was required to inhibit *Striga* seed germination than to inhibit the tillering suppression activity of GR24. This weak inhibition of *Striga* seed germination by KK094 is consistent with the difference in IC_50_ values for the hydrolysis activity of ShHTL7 and D14. However, modification of KK094 might improve its inhibitory effect on the functions of *Striga* HTL proteins. Thus, KK094 and its derivatives have the potential to regulate plant growth and seed germination of parasitic plants both in laboratories and in fields, and KK052/KK073 and their derivatives are expected to be powerful tools to elucidate the complicated mechanisms underlying SL-perception and signal transduction.

### Materials and Methods

#### Plant materials and the rice tillering assay

An SL-deficient rice mutant, *d10-2*, of the Japonica-type cultivar (*Oryza sativa* L. cv. Nipponbare) and a *d17-1* rice mutant of the Japonica-type cultivar (*Oryza sativa* L. cv. Shiokari) were used in this assay. The rice seeds were sterilized in a 2.5% sodium hypochlorite solution containing 0.02% Tween 20 for 20 min. The seeds were washed five times with sterilized water and then incubated in tubes filled with water at 25 °C in the dark for 2 days. The germinated seeds were planted in a hydroponic culture medium (*34*) solidified with 0.6% agar and cultivated at 25 °C under fluorescent light (70–100 µmol^−2^ sec^−1^) with a 16-hr light/8-hr dark photoperiod for 7 days. Each seedling was transplanted to a glass vial filled with 12 mL of sterilized hydroponic culture solution with or without an experimental compound and grown under the same conditions for 7 days.

#### Yeast two-hybrid (Y2H) and yeast three-hybrid (Y3H) assays

The Matchmaker Two-Hybrid System (Takara Bio, Otsu, Japan) was used for the Y2H assay. We used pGBK-D14 (*19*) as the bait and pGAD-SLR1 (*19*) or pGAD-D53 (*6*) as the prey. The *Saccharomyces cerevisiae* AH109 strain was transformed with the bait and prey plasmids and grown in liquid medium for 2 days. The plate assays (synthetically defined medium without histidine and adenine) were performed according to the manufacturer’s protocol, except that the plate medium contained various combinations of SLs and test compounds. For the Y3H assay, the yeast strains AH109 and the plasmids pGADT7 were also used and pBridge were obtained from Takara Bio Inc. pBridge-BD:D14-M:OSK1 was constructed by fusing D14 cDNA with the GAL4-BD domain and inserting OSK1 cDNA (Os11g0456300) into the site downstream of pMET25. pGAD-D3 was constructed by fusing D3 cDNA (AK065478) with the GAL-AD domain. AH109 was transformed with the pBridge-BD:D14-M:OSK1 and pGAD-D3 or pGADT7 selected on SD media lacking L-tryptophan (SD-Trp Leu Met). For the assay, the transformants were incubated on SD-Trp, Leu and Met media that lacked Ade and His (SD-His Ade Met).

#### *Striga* germination assay

*Striga* seed germination assay was performed as described previously (*35*). Seeds of *Striga hermonthica* harvested in Sudan were kindly provided by Professor A.E. Babiker (Sudan University of Science and Technology) and imported with the permission of the Minister of Agriculture, Forestry and Fisheries of Japan. Seeds of *S. hermonthica* were sterilized with a 1% sodium hypochlorite solution containing 0.01% Tween 20 for 5 min and washed five times with sterilized water. The seeds were then added to a 0.1% agar solution and dropped onto small, round, glass-fiber filters. Filters containing the seeds were arranged on a filter paper (70 mm diameter) in a Petri dish, and 1400 µL of sterilized water including the appropriate chemical was added to the dish. The dishes were incubated at 30 °C in the dark for 4 days. The small filters containing the seeds were transferred to a 96-well plate, and 10 µL of sterilized water or water including the appropriate chemical was added to each well. After incubation for 2 days under the same conditions, the number of germinated seeds was counted.

#### Protein preparation

D14 was expressed in *E. coli* and purified as described previously (*2*). Briefly, D14 (residues 54−318) from rice was expressed with the pET-49b expression vector (Merck-Millipore) in *E. coli* Rosetta (DE3) cells (Merck-Millipore). The cells were harvested, resuspended in extraction buffer (20 mM Tris-HCl (pH 8.5), 500 mM NaCl, 10% glycerol and 3 mM dithiothreitol (DTT)), and disrupted by sonication. The soluble fraction separated by centrifugation was purified using Glutathione Sepharose 4B resin (GE Healthcare). For crystallization, on-column cleavage was performed by adding HRV3C protease, and the eluted D14 was further purified using a Resource S column (GE Healthcare). The purified D14 was concentrated to 6.0 mg mL^−1^ in buffer containing 20 mM MES-NaOH (pH 6.5), 300 mM NaCl, 10% glycerol, and 5 mM DTT. For pull-down assays, GST fusion D14 was eluted with elution buffer (20 mM Tris-HCl (pH 7.8), 500 mM NaCl, 100 mM reduced glutathione and 5 mM DTT. After the eluted product was concentrated and diluted 25-fold in elution buffer lacking NaCl and reduced glutathione, GST-D14 was further purified using a Resource S column. The purified GST-D14 was concentrated to 5 μM in buffer containing 20 mM Tris-HCl (pH 7.8), 100 mM NaCl and 5 mM DTT. The *D53* open reading frame fragment was amplified by PCR using total complementary DNA from rice seedlings. For expression in *E. coli*, the PCR product was cloned into the expression vector pGEX-6P-3 (GE Healthcare), and subsequently transformed into *E. coli* Rosetta (DE3) cells. The cells were grown in Luria-Bertani broth at 37 °C to an OD_600_ of ~0.6 and induced with 0.5 mM isopropyl-β-_D_-thiogalactopyranoside at 15 °C for 32 h. The cells were harvested, resuspended in extraction buffer (20 mM Tris-HCl (pH 8.5), 500 mM NaCl and 5 mM DTT), and disrupted by sonication. The soluble fraction separated by centrifugation was purified using Glutathione Sepharose 4B resin. On-column cleavage was performed by adding HRV3C protease, and the eluted D53 was further purified using a Mono Q column and Superdex 200 columns (GE Healthcare). The purified D53 was concentrated to 5 μM in buffer containing 20 mM Tris-HCl (pH 7.8), 100 mM NaCl and 5 mM DTT for pull-down assays.

#### Hydrolysis assay

GR24 and KK094 pre-incubated at 30 °C for 10 min and incubated further at 30 °C for 20 min after the addition of 0.33 μM of D14. After adding 5 nmol of CPMF (5-(4-chlorophenoxy)-3-methylfuran-2(5*H*)-one) (36) as an internal standard, reaction solutions were extracted with ethyl acetate three times. Ethyl acetate layers were combined, dried in vacuo, and then dissolved with 50 μL methanol. Each sample was injected at a volume of 5 µl into the reverse-phase HPLC column (CAPCELL CORE C18, 2.1 Å ~100 mm, Shiseido, Tokyo, Japan), coupled to the ESI-MS system (LC-2030C/3D, Shimadzu, Kyoto, Japan). The analytes were eluted with a linear gradient of 40–90% buffer B (methanol with 0.1% (v/v) formic acid) in buffer A (MilliQ water with 0.1% (v/v) formic acid) within 9 min, keeping the final condition for 6 min. The column was operated at 40 °C with a flow rate of 0.2 mL min^−1^. Quantities of GR24 or ABC-OH were monitored at m/z = 299.10 or 202.06, respectively, and normalized by the amount of CPMF monitored at m/z = 224.02. The amount of GR24 incubated without D14 was set to 1. Error bars indicate SE of six seedlings. Student’s *t*-test was used to determine the significance of differences (\**p*<0.05, \*\**p*<0.01).

#### Yoshimulactone G *in vitro* analysis

In the hydrolysis assay, 0.3 μM of YLG was reacted with 0.33 μM of recombinant proteins (D14 or ShKAI2d6/ShHTL7) in a reaction buffer (100 mM PBS buffer, pH 7.3) with 0.1% dimethyl sulfoxide (DMSO) at a 300 μL volume on a 96-well black plate (Thermo). The fluorescent intensity was measured by ThermoFisher at the excitation by 485 nm, and detected at a wavelength of 535 nm. The enzymatic reaction was carried out in a 30 °C incubator for 15 min. IC_50_ values were calculated using the website: http://www.ic50.tk/index.html.

#### Differential Scanning Fluorometry

DSF experiments were performed using CFX Connect Real-Time PCR Detection System (Bio-Rad, CA). Sypro Orange was used as a reporter dye. After pre-incubation of the reaction mixtures for 5 min and 10 s at 20 °C, reaction mixtures were denatured using a linear 20 °C to 95 °C gradient at a rate of 0.5 °C per 10 s in the absence of light. Each reaction was carried out at 20-μL scale in TN buffer (10 mM Tris-HCl, pH=8.0, 200mM NaCl) containing 5 μM protein, 0.01 μL Sypro Orange, and each concentration of experimental compounds (final DMSO concentration was 10%). The experiments were repeated twice.

#### Pull-down assays

Pull-down assays of GST-D14 and D53 were performed using a two-step treatment with experimental compounds. The purified GST-D14 (5 μM) was incubated at 4 °C for 30 min with Glutathione Sepharose 4B resin, and then washed three times. The glutathione agarose-bound GST-D14 was incubated with or without 50 μM of each experimental compound at room temperature for 2 h (first step) and then washed three times. The resin-bound GST-D14 was incubated with D53 (5 μM) with or without 50 μM of each experimental compound at room temperature for 2 h (second step), then washed three times. Eluted proteins were separated using SDS-PAGE electrophoresis (10% gel) and detected using CBB staining.

#### Crystallization and structure determination

6.0 mg/ml of D14 protein and 10 mM KK094/KK052/KK073 were mixed together and subjected to crystallization by the sitting drop vapor diffusion method. Crystals of the D14–KK094CM complex were obtained at 20 °C using a reservoir solution containing 100 mM MES (pH 6.5) and 13% PEG20000. Crystals of the D14– KK052CM complex were obtained at 20 °C using a reservoir solution containing 100 mM HEPES (pH 7.5) and 13% PEG20000. Crystals of the D14–KK073CM complex were obtained at 20 °C using a reservoir solution containing 100 mM MES (pH 6.5) and 11% PEG 20000. All crystals were soaked in the cryo-protectant solution containing 25% (v/v) ethylene glycol and then flash-cooled with a nitrogen-gas stream at 100 K. All X-ray diffraction data were collected on the BL-1A beamline at the Photon Factory (Tsukuba, Japan) and processed using the XDS package (*37*). Molecular replacement was performed using Phaser (*38*) in PHENIX (*39*) with the apo-D14 structure (PDB ID 3VXK) (*19*) as the initial model. Coot (*40*) was used to manually fit the protein models, and the structure was refined with PHENIX. The geometry of the final model was analyzed using RAMPAGE (41), and superposition and r.m.s.d. of the structures were calculated using the CCP4 program LSQKAB (*42*). All structure Figures were prepared using PyMOL (*43,44*). X-ray data and refinement statistics are given in table S1. Coordinates of the X-ray structures of the D14–KK094CM complex, the D14–KK052CM complex, and the D14– KK073CM complex have been deposited in the Protein Data Bank, under accession codes 5ZHR, 5ZHS and 5ZHT, respectively.

#### Mass Spectrometry

For the identification of the KK094CM binding site, the reaction mixtures (ca. 1.6 mg mL^−1^) of D14 with or without KK094 were dialyzed against 50 mM of NH_4_HCO_3_ solution. Aliquots of the dialyzed samples were analyzed using MALDI-TOF MS on an ultrafleXtreme TOF/TOF Mass Spectrometer (Bruker Daltonics, Bremen, Germany) in a linear mode using sinapinic acid as a matrix. After confirmation of KK094 binding to the protein using MALDI-TOF MS, the dialyzed samples were digested with trypsin (TPCK treated, Worthington Biochemical Co.) overnight at 37 °C. Aliquots of the digestion mixture were also applied to MALDI-TOF MS in reflector mode using α-cyano-4-hydroxycinnamic acid as a matrix. For binding site determination, the digests were further analyzed by nano-liquid chromatography-tandem mass spectrometry using a Q Exactive mass spectrometer (Thermo Fisher Scientific). The peptide mixtures (2 μL) were separated using a nano ESI spray column (75 μm [ID] × 100 mm [L], NTCC analytical column C18, 3 μm, Nikkyo Technos, Tokyo, Japan) with a linear gradient of 0–35% buffer B (acetonitrile with 0.1% (v/v) formic acid) in buffer A (MilliQ water with 0.1% (v/v) formic acid) at a flow rate of 300 nL min^−1^ over 10 min (EAST-nLC 1000; Thermo Fisher Scientific). The mass spectrometer was operated in the positive-ion mode, and the MS and MS/MS spectra were acquired using a data-dependent TOP10 method. The MS/MS spectra were drawn using the Qual Browser, Thermo Xcalibur 3.1.66.10.

#### Preparation of KK compounds

A description is available in the **Supplementary Notes**.

## Acknowledgments

The authors thank Prof. A.E. Babiker (Sudan University of Science and Technology) for providing the seeds of *S. hermontica* harvested in Sudan and thank Prof. J. Kyozuka (Tohoku University) for kindly providing rice SL mutants.

## Author contributions

H.N., M.T., and T.A. designed research; H.N. and W.H. performed the Y2H assay, inhibitor screening and *in planta* assays and the *in vitro* enzymatic assays. K.H., T.M., and Y.X. performed the crystal structure analysis and pull-down assay; K.J. performed the DSF assay; N.D. performed the MALDI-TOF-MS and LC-MS/MS analyses; K.K. synthesized chemicals; H.N., K.H., and T.A. wrote the paper; all authors read and approved the manuscript.

## Data and materials availability

The structure coordinates and structural factors are deposited in the Protein Data Bank with accession numbers of 5ZHR (D14–KK094CM complex), 5ZHS (D14–KK052CM complex) and 5ZHT (D14−KK073CM complex).

## Supplementary Materials

**fig. S1.**
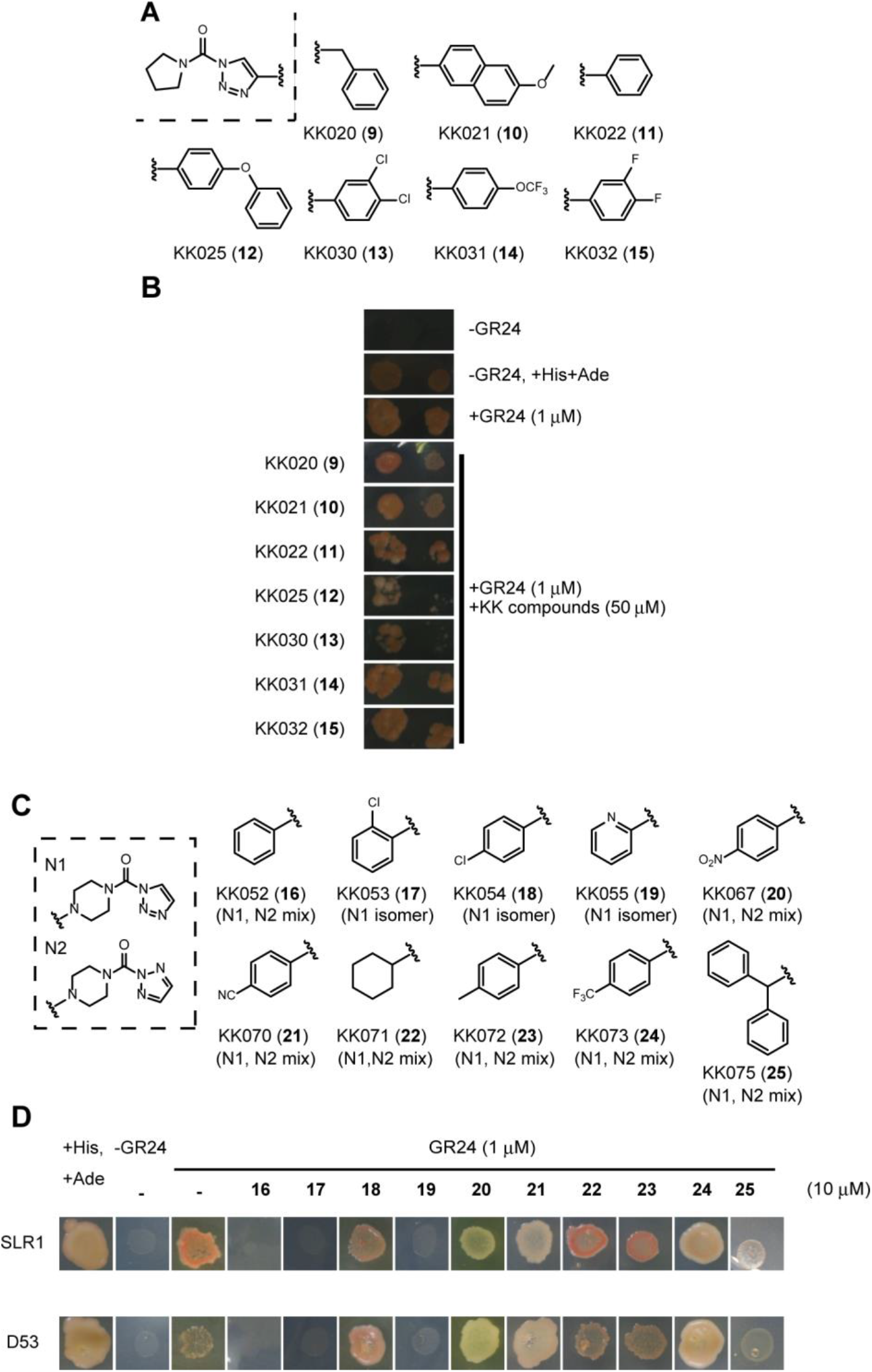
Inhibition of the SL-dependent D14–D53 interaction by KK007-and KK004-derivatives. (**A**) Structures of KK007N1-derivatives. Various substituents were introduced onto the 4-position of the 1,2,3-triazole group of KK007N1. (**B**) Growth of AH109/pGBK-D14–pGAD-D53 on SD-His Ade plates containing 1 μM GR24 with 10 μM or 50 μM 1,2,3-triazole ureas for 4 days at 30 °C. (**C**) Structures of KK004-derivatives. The morphorine structure derived from KK004 (mixture of N1- and N2-compounds; shown in the dashed box) was modified by the addition of various substituents. (**D**) Growth of AH109/pGBK-D14–pGAD-D53 on an SD-His Ade plate containing 1 μM GR24 with 10 μM or 10 μM of 1,2,3-triazole ureas for 4 days at 30 °C.

**fig. S2.**
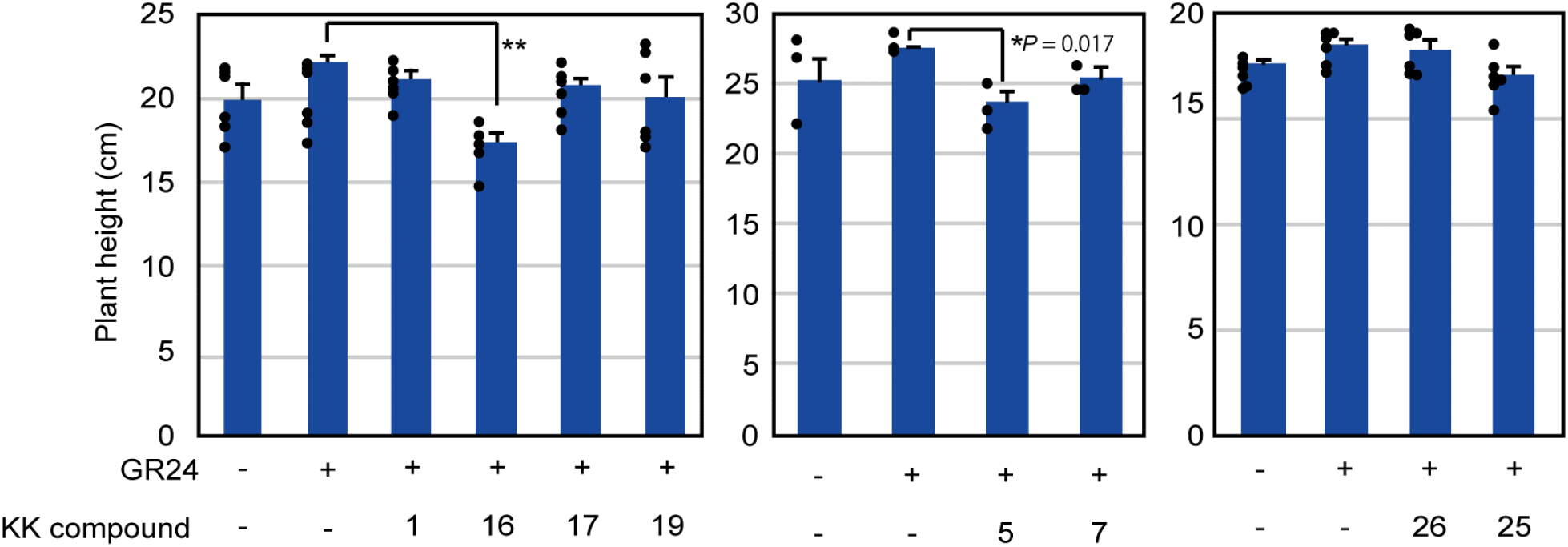
Effects of KK compounds on plant growth. Seven-day-old *d17-1* mutant rice seedlings were treated with experimental compounds, grown for a further 7 days, then plant heights were measured. Error bars indicate SE of six seedlings. Unpaired tw-Student’s *t*-test was used to determine the significance of differences (\**p*<0.05, \*\**p*<0.01).

**fig. S3.**
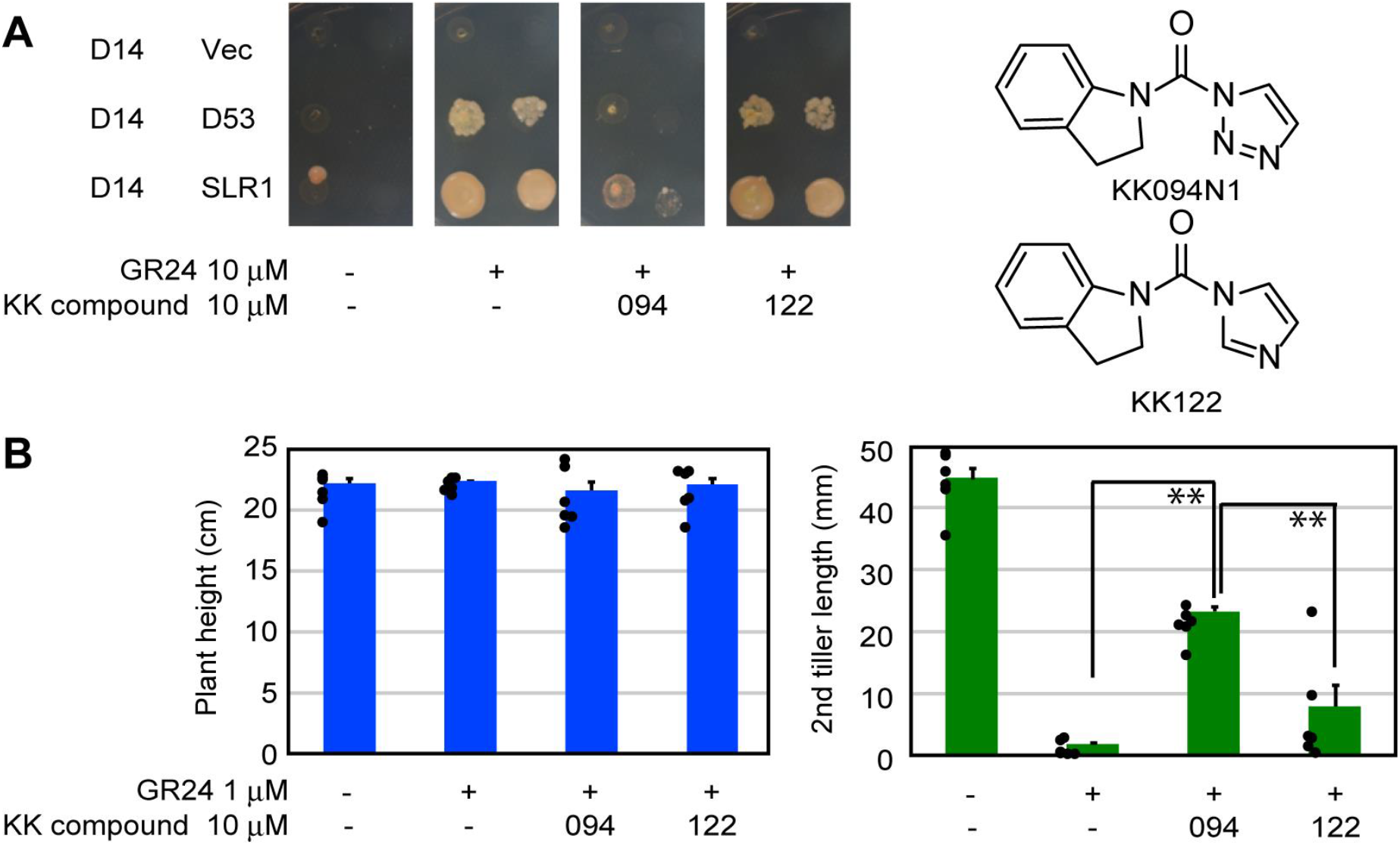
Inhibition of the SL-dependent D14–D53 interactions and rice tillering by KK122. (**A**) Growth of AH109 yeast transformed with pGBK-D14 and pGAD-T7/pGAD-D53/pGAD-SLR1 on an SD-His Ade plate containing 10 μM GR24 with 10 μM KK094N1/KK122 for 4 days at 30 °C. (**B**) Inhibition of the effect of SL on rice tillering by KK122. Seven-day-old *d17-1* mutant rice seedlings were treated with experimental compounds and grown for a further 7 days, and plant heights and the length of the first/second tillers were measured. Error bars indicate SE of six seedlings. Unpaired two-tailed Student’s *t*-test was used to determine the significance of differences (\*\**p*<0.01).

**fig. S4.**
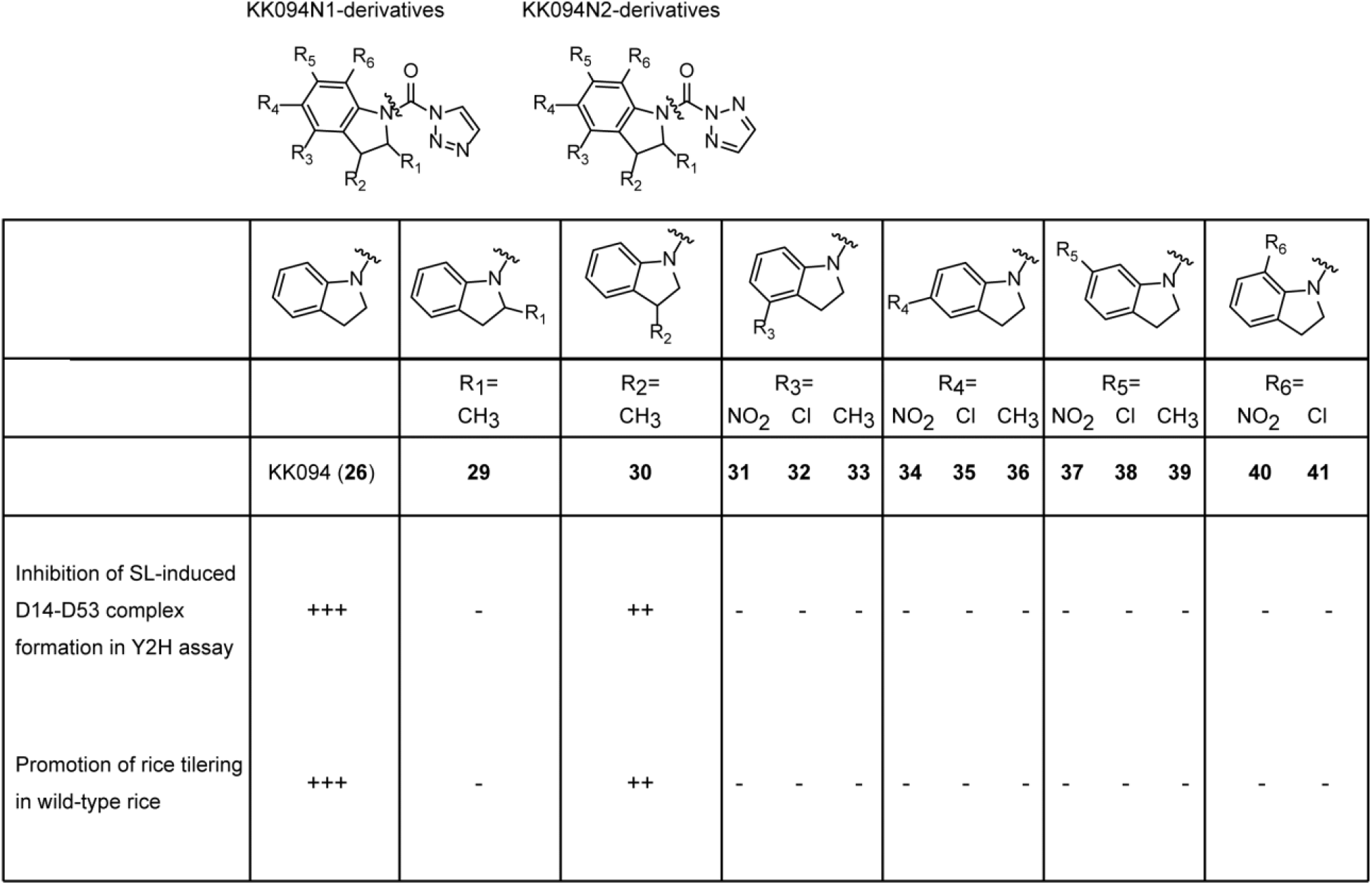
Inhibition of the SL-dependent D14–D53 interactions and rice tillering by KK094-derivatives. Upper line in the bottom column: Growth of AH109 yeast transformed with pGBK-D14 and pGAD-T7/pGAD-D53 on an SD-His Ade plate containing 1 μM GR24 with experimental compounds for 4 days at 30° C. The extent of growth inhibition is indicated by – to +++ (- no inhibition; ++ partial inhibition; +++ complete inhibition). Lower line in the bottom column: Inhibition of the effect of SL on rice tillering. Seven-day-old *d17-1* mutant rice seedlings were treated with/without 1 μM GR24 and 10 μM KK094-derivatives and grown for 7 days before the length of second tillers were measured. The extent of tillering bud growth promotion is presented by – to +++ (- no promotion [2nd tiller length = 0 cm]; + weak promotion [2nd tiller length = 0–1.0 cm]; ++ intermediate inhibition [2nd tiller length = 1.0–3.0 cm]; +++ strong promotion [2nd tiller length > 3.0 cm]).

**fig. S5.**
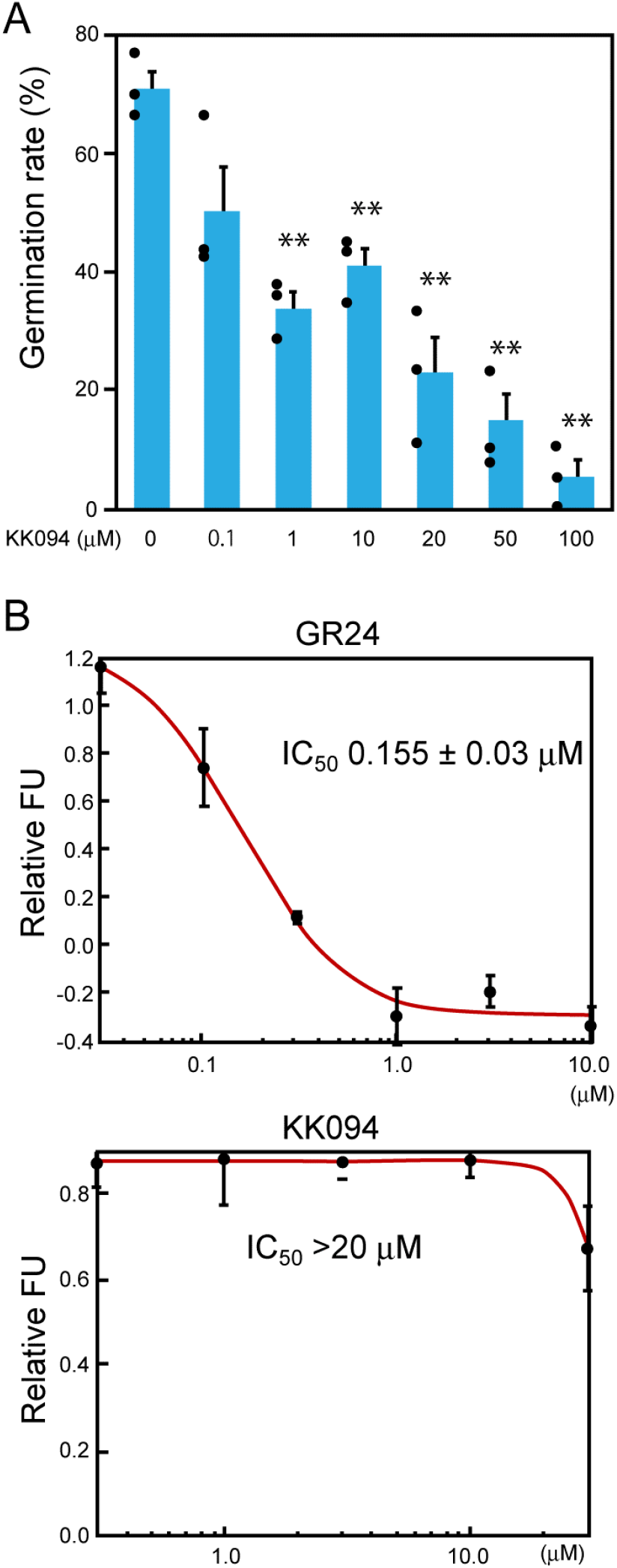
Inhibition of *Striga* seed germination and a *Striga* SL receptor, ShHTL7, by KK094. (**A**) KK094 concentration-dependently inhibits seed germination of *Striga hermonthica*. Germination ratios of *Striga* seeds treated with 1 μM of GR24 and KK094 at the concentrations shown. Error bars indicate SE of three independent experiments. Unpaired two-tailed Student’s *t-*test was used to determine the significance of differences with the germination rate of KK094-non-treated seeds (\*\**p*<0.01). (**B**) Competitive inhibition of ShHTL7-mediated YLG hydrolysis. Following reaction of 0.3 μM of YLG with 0.33 μM of D14, the fluorescent intensity was measured at the excitation wavelength of 485 nm and detected at a wavelength of 535 nm. The fluorescent values relative to the hydrolysis without inhibitor is shown. FU = fluorescence unit. IC50 values were calculated using the website: http://www.ic50.tk/index.html.

**fig. S6.**
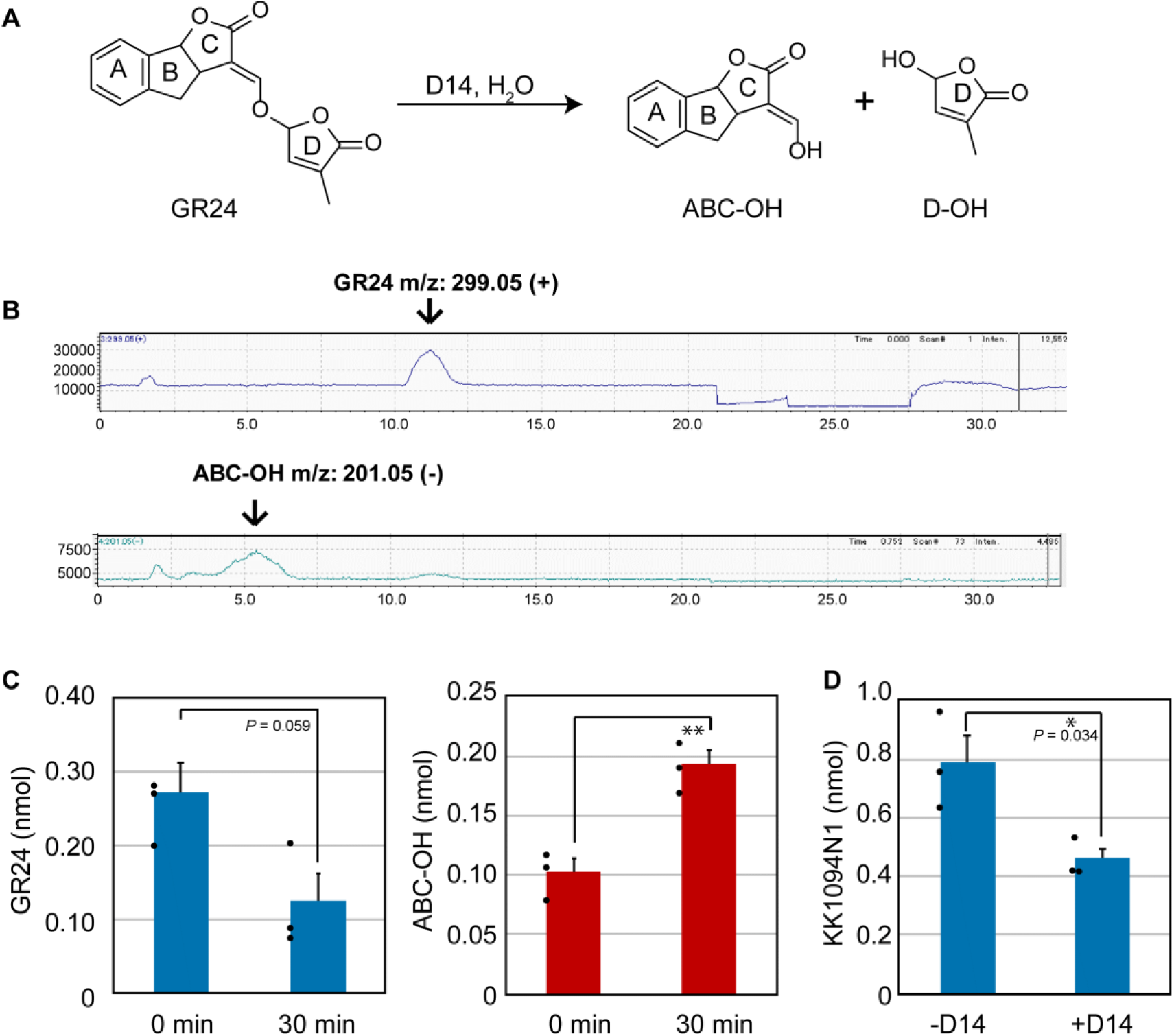
LC-MS analyses of GR24- or KK094N1-hydrolysis by D14. (**A**) The reaction formula of D14-mediated hydrolysis of GR24. (**B–D**) GR24 or KK094 was pre-incubated at 30 °C for 10 min and incubated further at 30 °C for 20 min after addition of 0.33 μM of D14. Then quantities of GR24 or ABC-OH were monitored using LC-MS (**B**, **C**). Quantities of KK094N1 were also monitored by LC-MS (**D**). Error bars indicate SE of three sample replicates. Unpaired two-tailed Student’s *t*-test was used to determine the significance of differences (\**p*<0.05, \*\**p*<0.01).

**fig. S7.**
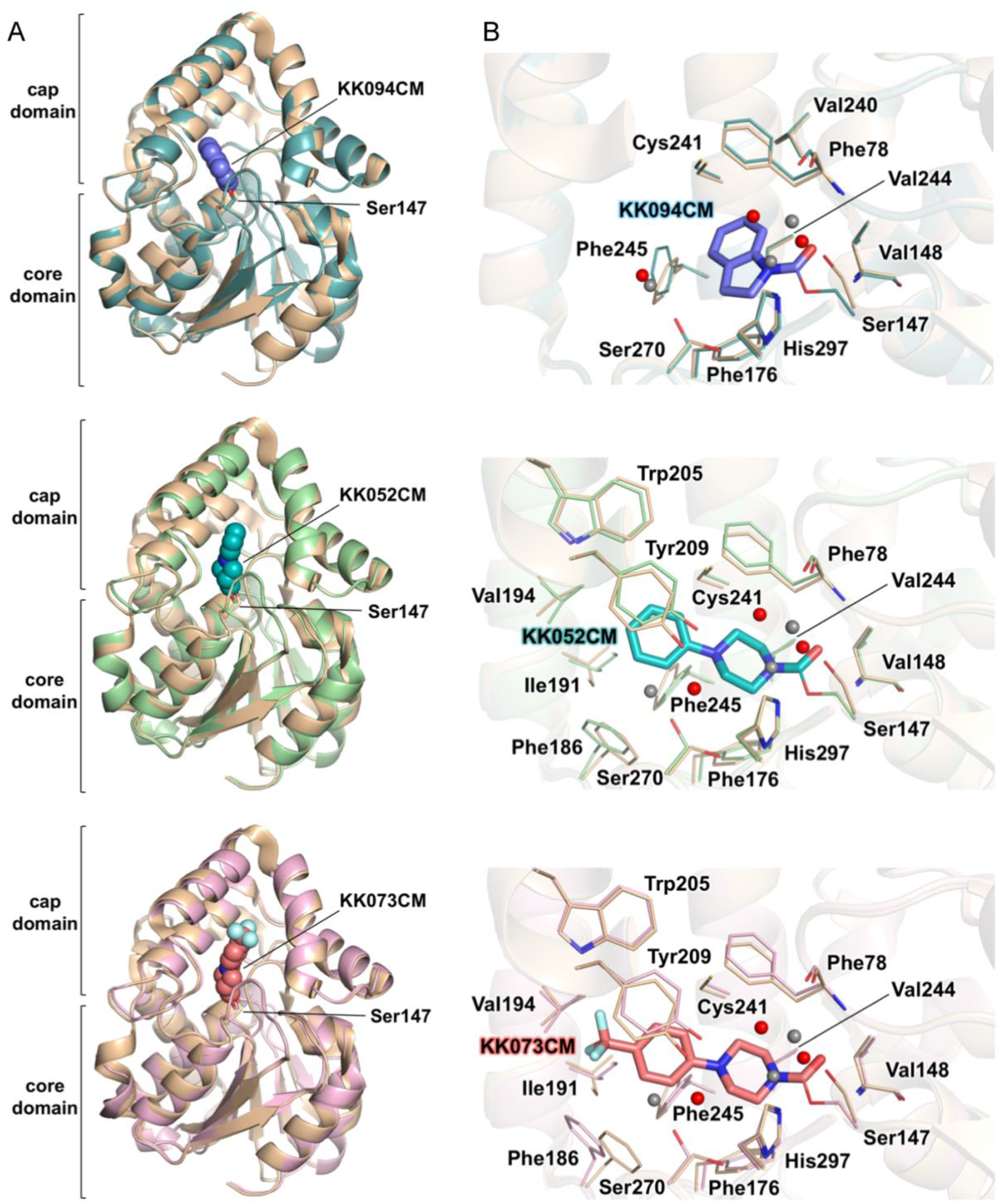
Structural comparisons. (**A**) KK094CM-bound D14, KK042CM-bound D14, and KK073CM-bound D14 are superimposed on apo-D14 (PDB ID 3VXK) (white). The orientation and color coding are the same as in Fig. 3**A** and Fig. 6**A**. Each r.m.s.d. for the Cα atoms was 0. 16 Å (KK094CM), 0.35 Å (KK052CM), and 0. 35 Å (KK073CM). (**B**) The close-up view of the ligand-binding sites in (**A**). The orientation and structural representation are the same as in Fig. 3**A** and 6**C**. Water molecules from apo-D14 are shown as white balls.

**fig. S8.**
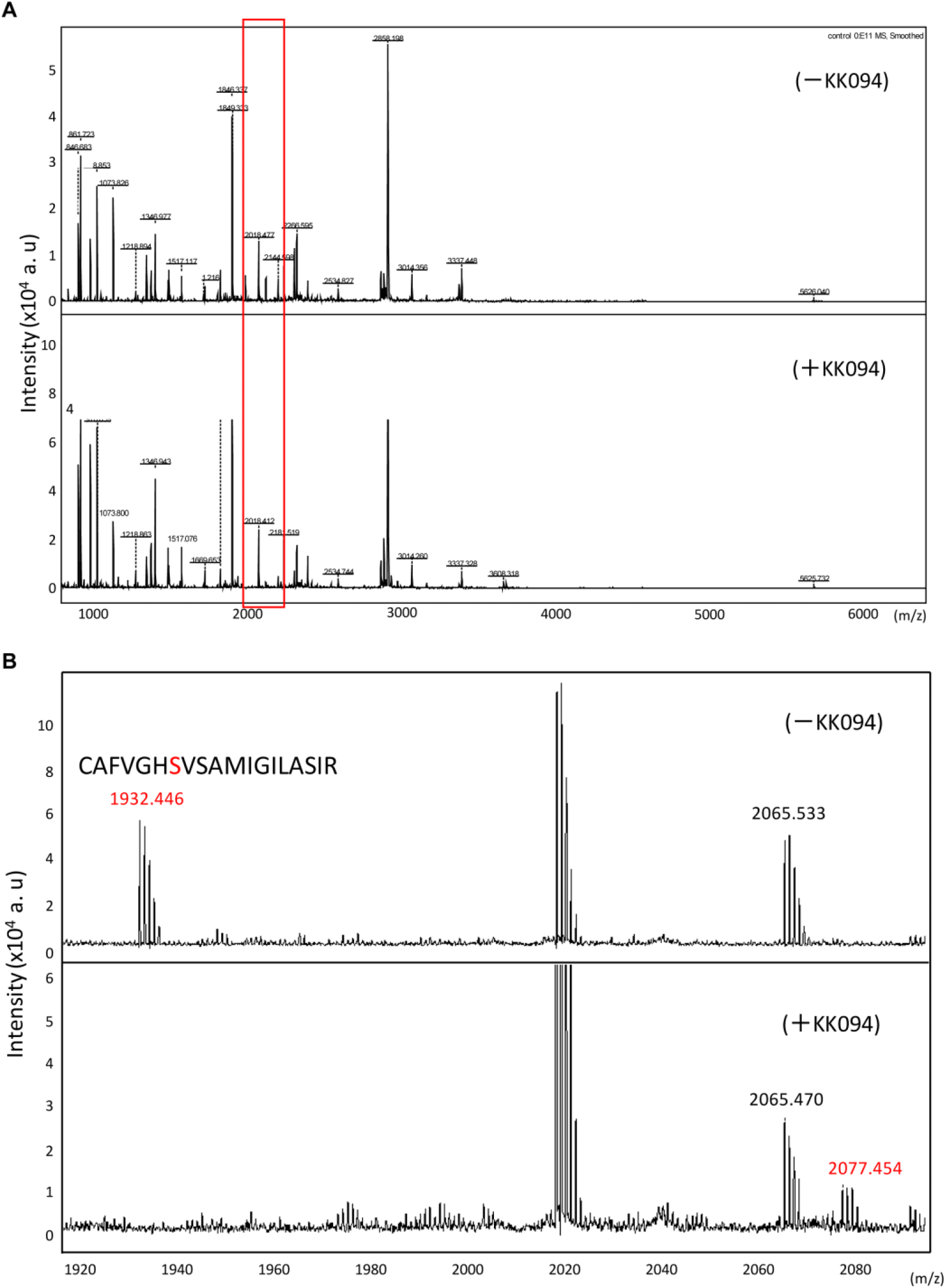
MALDI-TOF-MS analysis of trypsin-treated D14 with or without KK094. (**A**) D14 was incubated without (upper panel) or with (lower panel) KK094. The reaction mixture of D14 without or with KK094 (-KK094/+KK094, respectively) was dialyzed against 50 mM NH_4_HCO_3_ solution. Dialyzed sample was digested with trypsin and analyzed using MALDI-TOF-MS. (**B**) Magnification of the red box in panel (**A**). The fragment corresponding to the peptide 141–159 (^141^CAFVGHSVSAMIGILASIR^159^; *m*/*z* 1932.446) was detected under the -KK094 condition but was not detected under the +KK094 condition. On the contrary, the peptide fragment with a molecular mass of 2077.454, corresponding to the peptide 141–159 with covalently-bound KK094-CM was detected when D14 was incubated with KK094.

**fig. S9.**
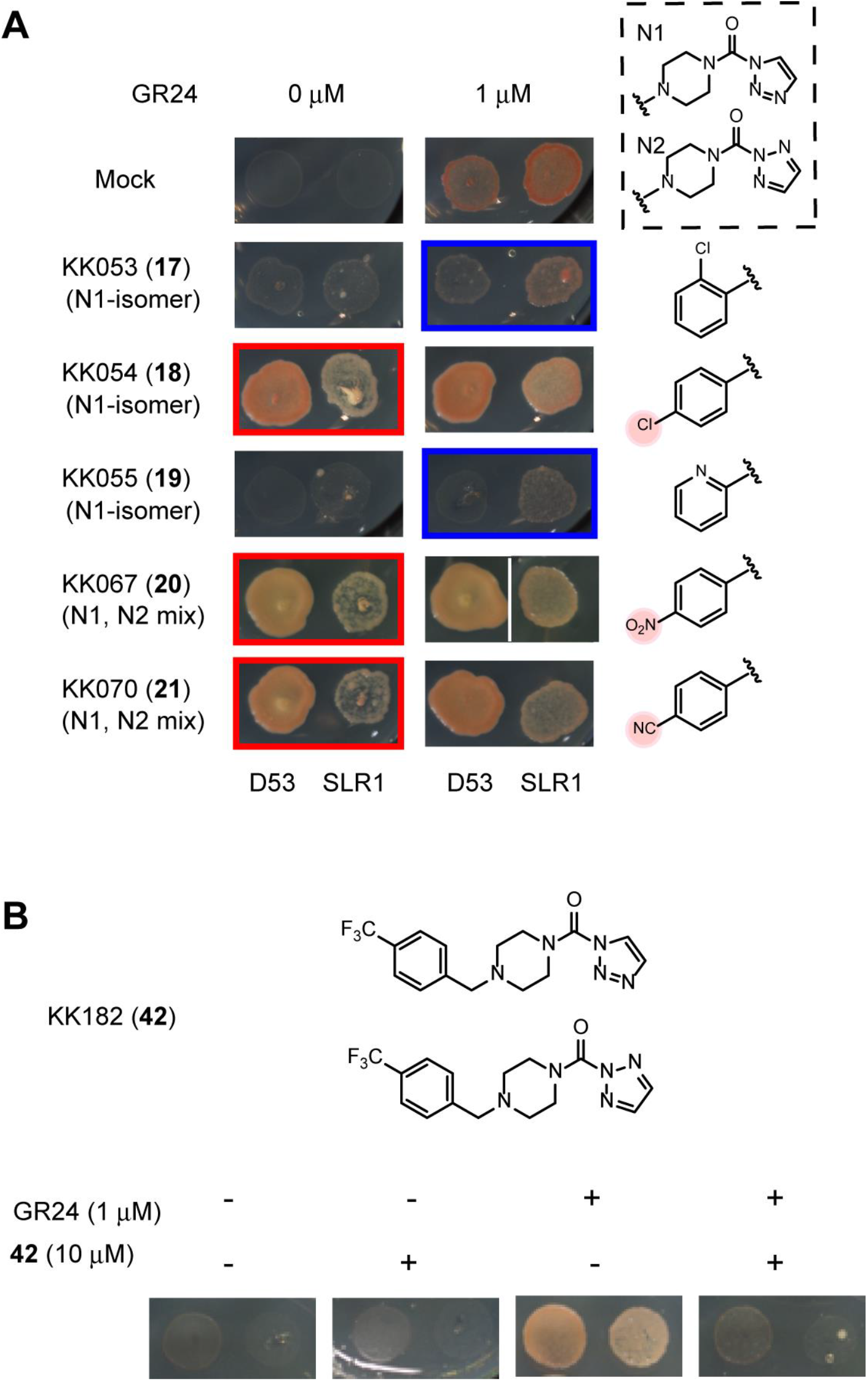
Inhibition or stimulation of the SL-dependent D14–SLR1/D14–D53 interaction by KK052-derivatives. (**A**) Structures of KK052-derivatives are shown on the right. The morphorine structure derived from KK052 (mixture of N1- and N2-compounds; shown in the dashed box) were modified by various substituents. Growth of AH109/pGBK-D14–pGAD-D53 or AH109/pGBK-D14–pGAD-SLR1 on an SD-His Ade plate containing experimental compounds for 4 days at 30 °C. Three compounds showed an agonist effect (in red squares), and two showed antagonism (in blue squares). (**B**) Structures of KK182 (a mixture of N1- and N2-compounds) and growth of AH109/pGBK-D14–pGAD-D53 on an SD-His Ade plate containing experimental compounds for 4 days at 30 °C.

**fig. S10.**
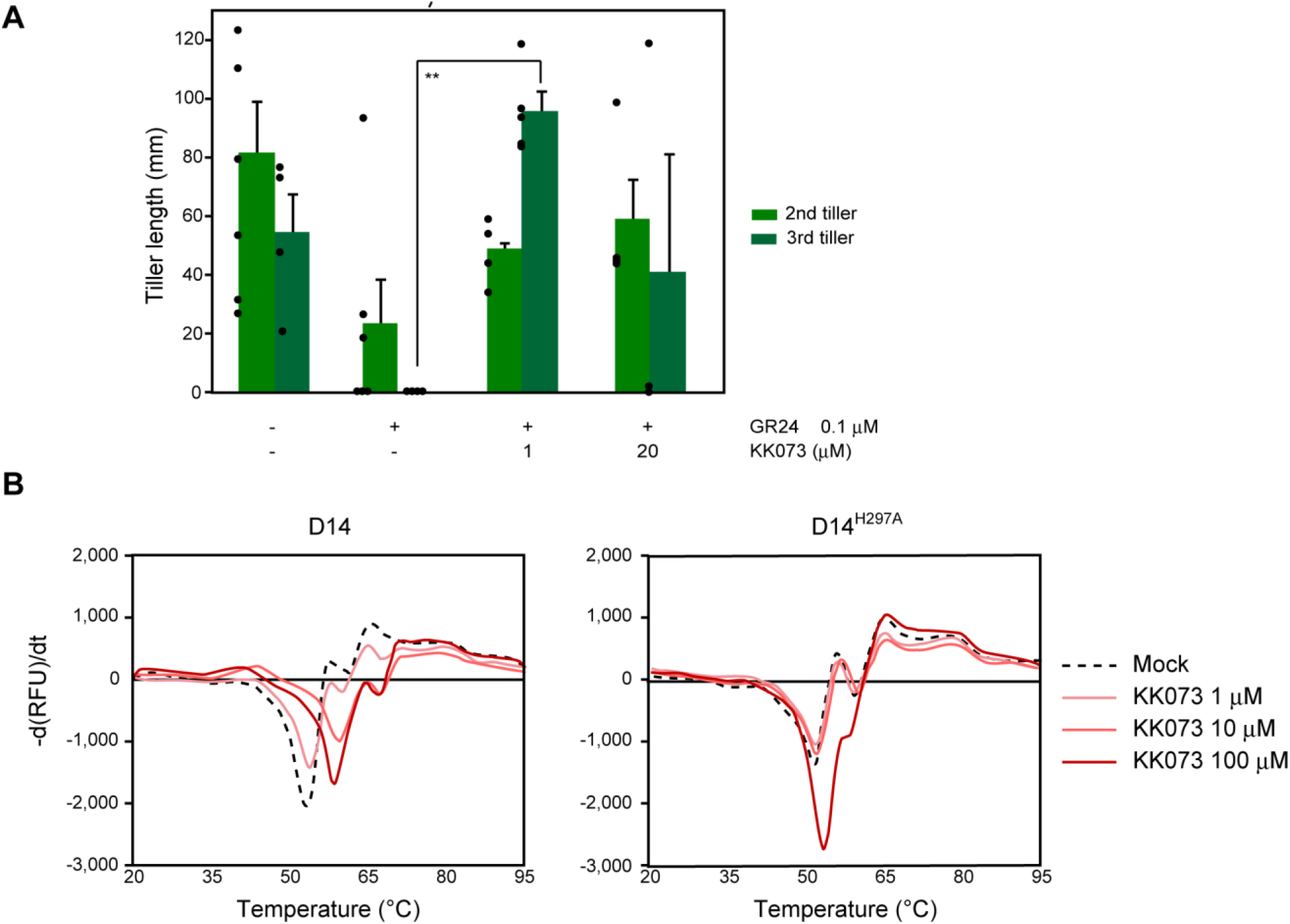
KK073 showed antagonism of the suppression of rice tillering induced by strigolactone. (**A**) KK073 showed antagonism of the suppression of rice tillering induced by strigolactone. (**A**) KK073 produced attenuation of the tillering inhibition induced by GR24. Seven-day-old *d10-1* mutant rice seedlings were treated with experimental compounds and grown for a further 2 weeks, then the length of second/third tillers were measured. Error bars indicate SE of three to six seedlings. Unpaired two-tailed student’s *t*-test was used to determine the significance of differences (\*\**p*<0.01). (**B**) Melting temperature curves for D14 and D14^H297A^ mutant proteins at varying concentrations of KK073 as monitored by differential scanning fluorimetry. Each line represents the average protein melt curve for three replicate samples measured in parallel.

**table S1.**
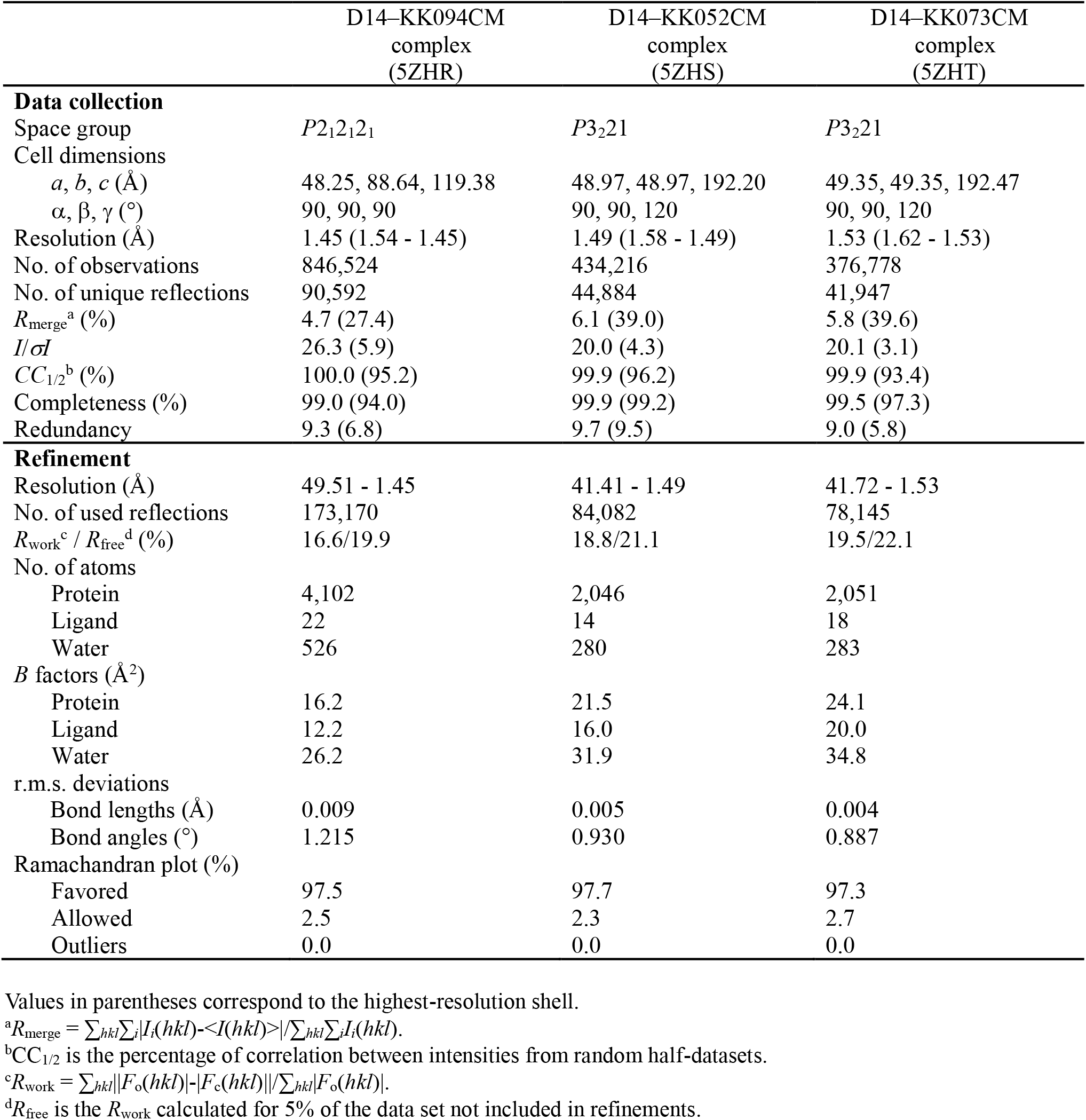
X-ray data collection and refinement statistics.

**table S2.**
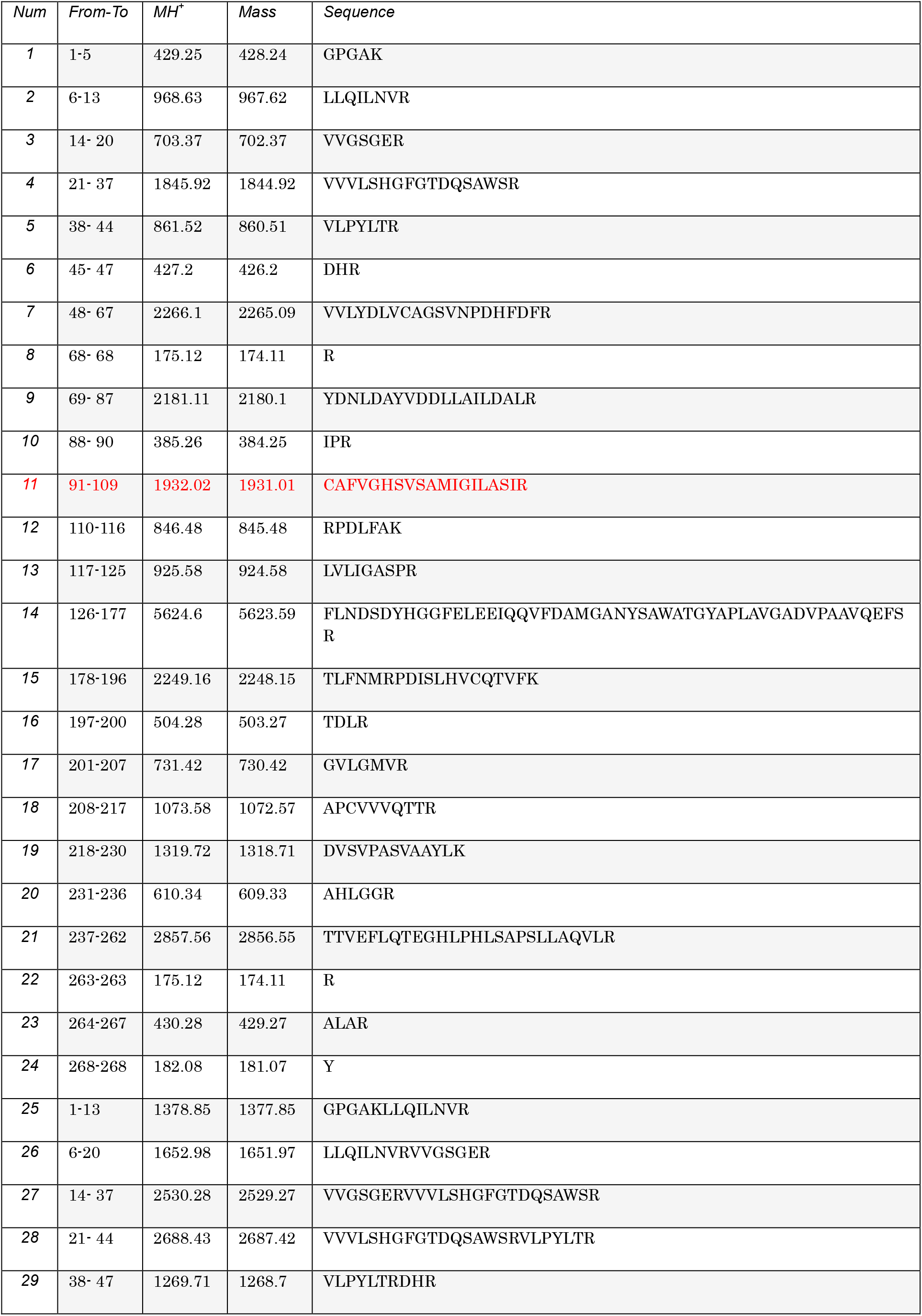
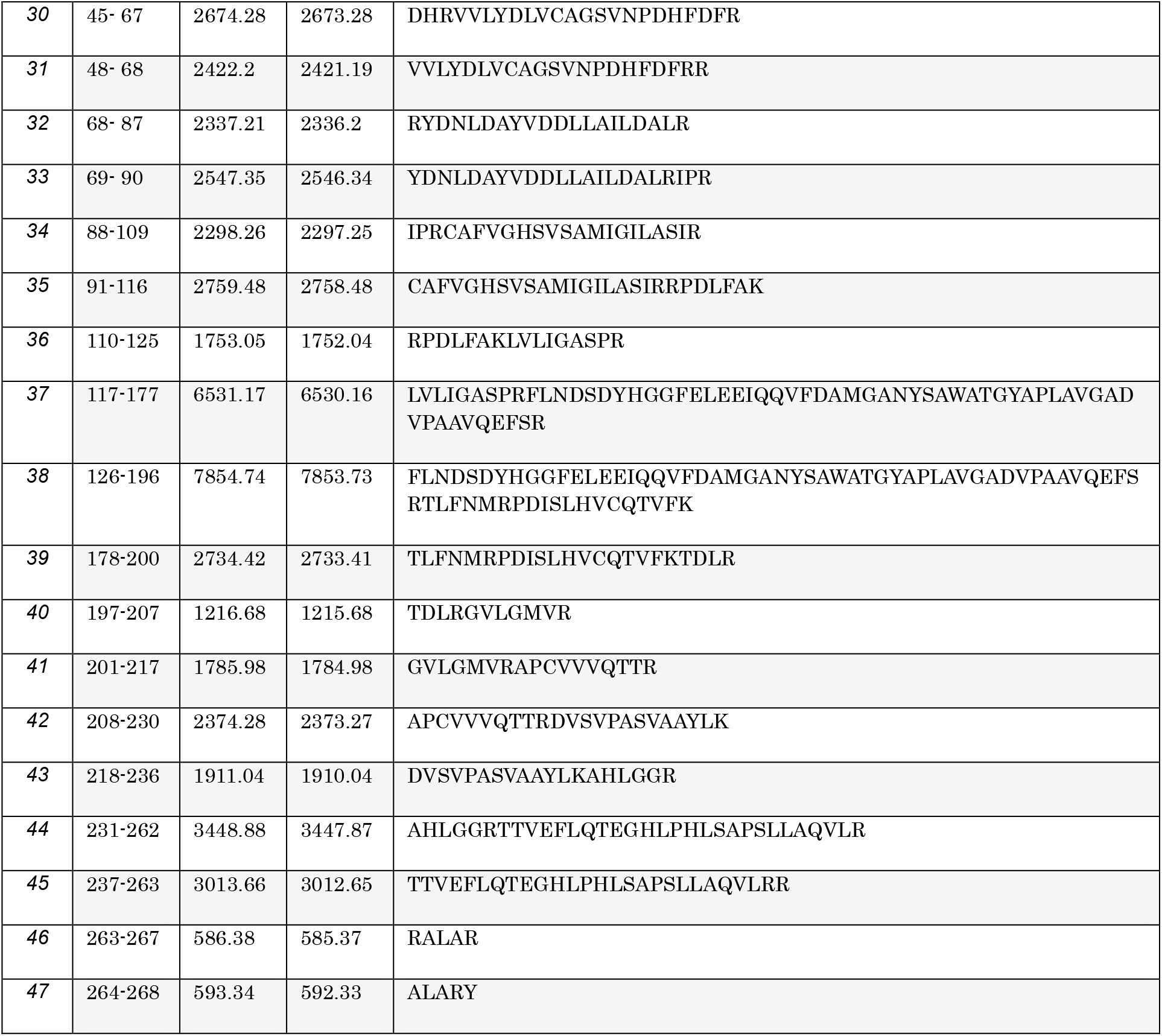
Deduced peptide mass values of D14 fragments digested with trypsin.

**table S3.**
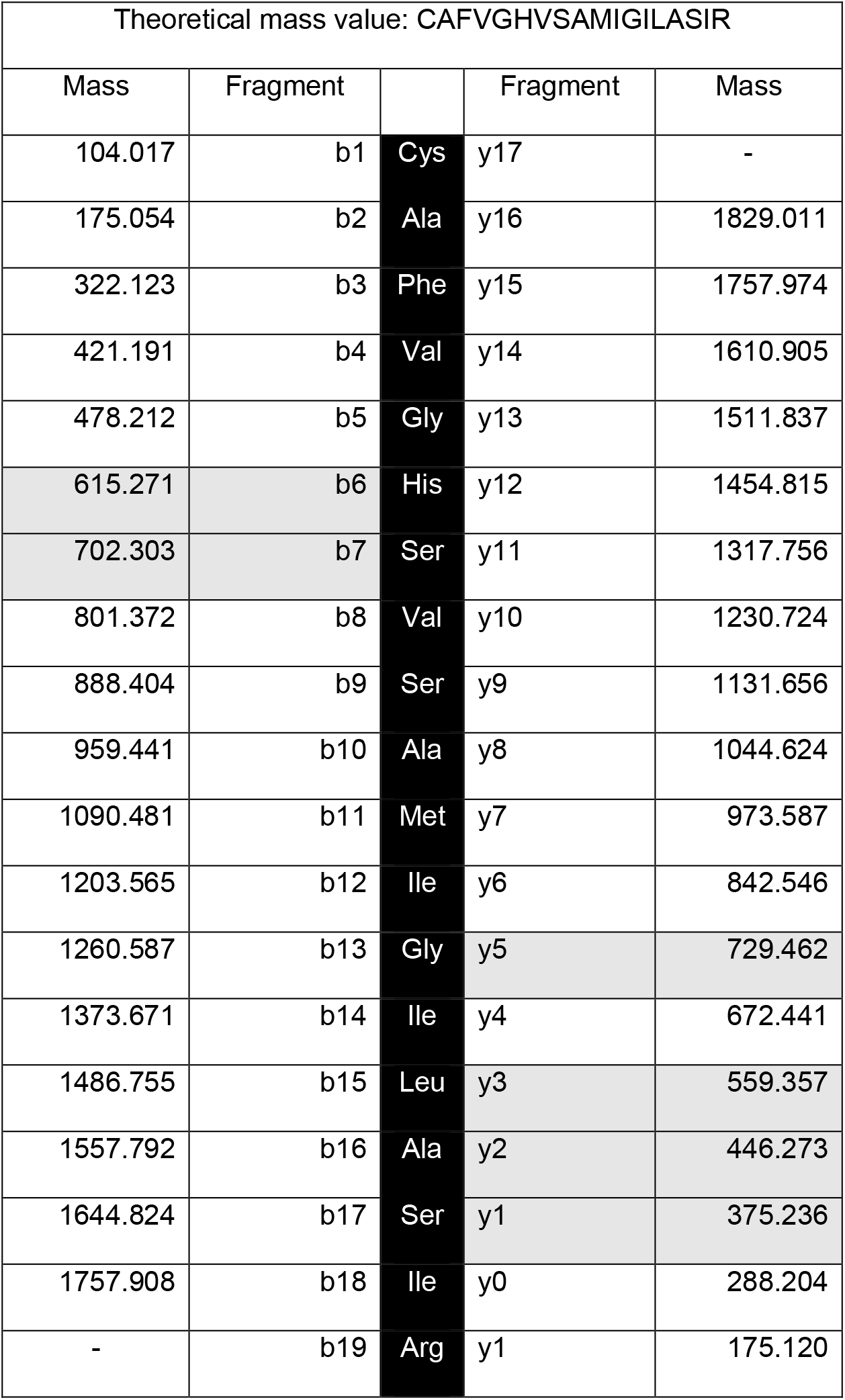
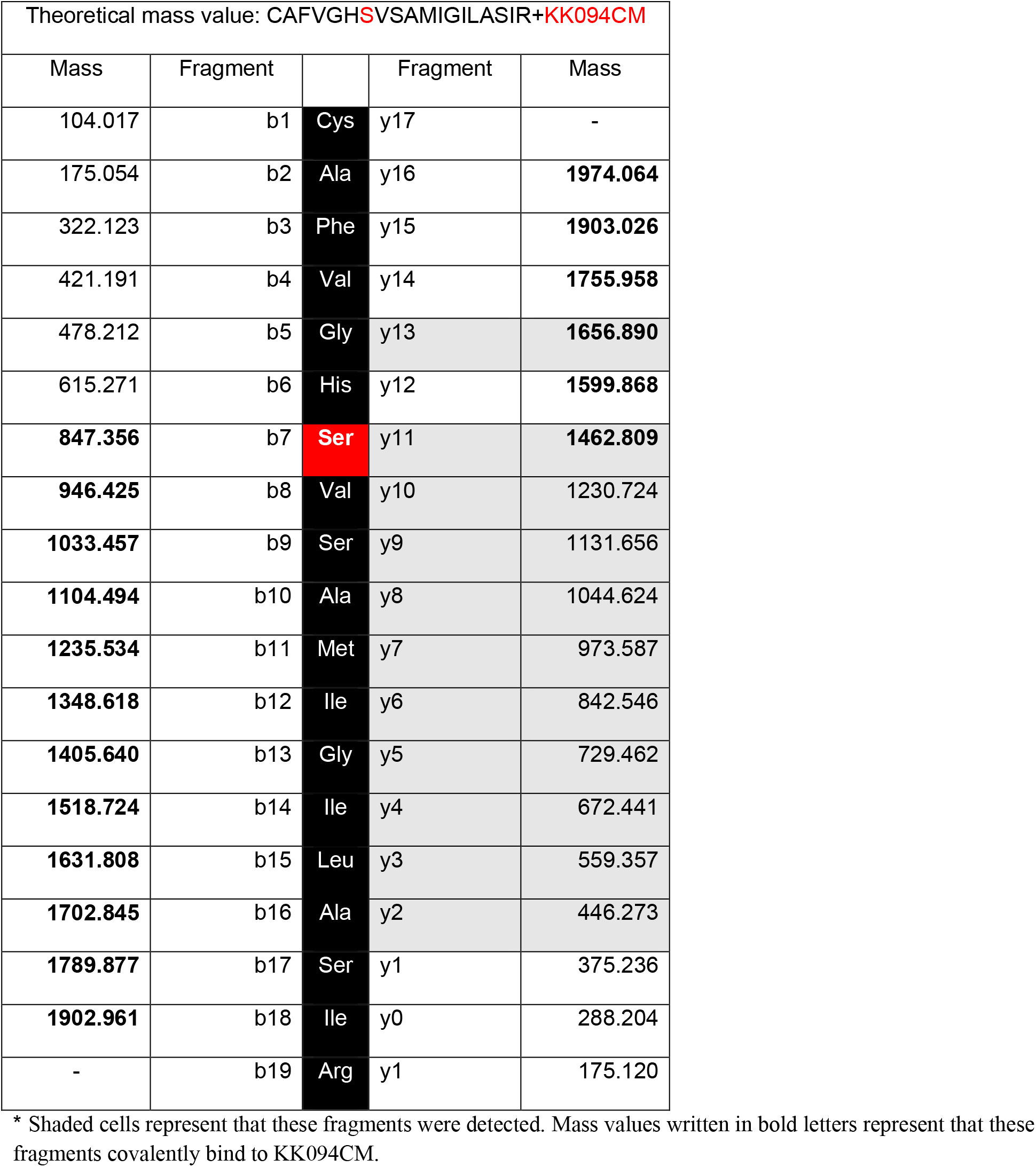
Theoritical mass values of peptide 141–159 (^141^CAFVGHSVSAMIGILASIR^159^).

| Theoretical mass value: CAFVGHSVSAMIGILASIR |  |  |  |  |
| --- | --- | --- | --- | --- |
| Mass | Fragment |  | Fragment | Mass |
| 104.017 | b1 | Cys | y17 | - |
| 175.054 | b2 | Ala | y16 | 1829.011 |
| 322.123 | b3 | Phe | y15 | 1757.974 |
| 421.191 | b4 | Val | y14 | 1610.905 |
| 478.212 | b5 | Gly | y13 | 1511.837 |
| 615.271 | b6 | His | y12 | 1454.815 |
| 702.303 | b7 | Ser | y11 | 1317.756 |
| 801.372 | b8 | Val | y10 | 1230.724 |
| 888.404 | b9 | Ser | y9 | 1131.656 |
| 959.441 | b10 | Ala | y8 | 1044.624 |
| 1090.481 | b11 | Met | y7 | 973.587 |
| 1203.565 | b12 | Ile | y6 | 842.546 |
| 1260.587 | b13 | Gly | y5 | 729.462 |
| 1373.671 | b14 | Ile | y4 | 672.441 |
| 1486.755 | b15 | Leu | y3 | 559.357 |
| 1557.792 | b16 | Ala | y2 | 446.273 |
| 1644.824 | b17 | Ser | y1 | 375.236 |
| 1757.908 | b18 | Ile | y0 | 288.204 |
| - | b19 | Arg | y1 | 175.120 |

| Theoretical mass value: CAFVGHSVSAMIGILASIR+KK094CM |  |  |  |  |
| --- | --- | --- | --- | --- |
| Mass | Fragment |  | Fragment | Mass |
| 104.017 | b1 | Cys | y17 | - |
| 175.054 | b2 | Ala | y16 | 1974.064 |
| 322.123 | b3 | Phe | y15 | 1903.026 |
| 421.191 | b4 | Val | y14 | 1755.958 |
| 478.212 | b5 | Gly | y13 | 1656.890 |
| 615.271 | b6 | His | y12 | 1599.868 |
| 847.356 | b7 | Ser | y11 | 1462.809 |
| 946.425 | b8 | Val | y10 | 1230.724 |
| 1033.457 | b9 | Ser | y9 | 1131.656 |
| 1104.494 | b10 | Ala | y8 | 1044.624 |
| 1235.534 | b11 | Met | y7 | 973.587 |
| 1348.618 | b12 | Ile | y6 | 842.546 |
| 1405.640 | b13 | Gly | y5 | 729.462 |
| 1518.724 | b14 | Ile | y4 | 672.441 |
| 1631.808 | b15 | Leu | y3 | 559.357 |
| 1702.845 | b16 | Ala | y2 | 446.273 |
| 1789.877 | b17 | Ser | y1 | 375.236 |
| 1902.961 | b18 | Ile | y0 | 288.204 |
| - | b19 | Arg | y1 | 175.120 |

\* Shaded cells represent that these fragments were detected. Mass values written in bold letters represent that these fragments covalently bind to KK094CM.

## Supplementary Notes

### General Procedure

NMR spectra (^1^H, ^13^C) were recorded at 500 MHz on a JNM-A500 spectrometer (JEOL, Tokyo, Japan) or a VNMR500 spectrometer (Agilent, CA). High resolution mass spectra (HRMS) were determined by electrospray ionization coupled to a time-of-flight analyser (Triple TOF 5600+ system, SCIEX, MA).

### KK002-N1 (1), KK002-N2 (2)

*N*,*N*-diethyl-1*H*-1,2,3-triazole-1-carboxamide (**1**), *N*,*N*-diethyl-2*H*-1,2,3-triazole-2-carboxamide (**2**).

The mixture of diethylcarbamoyl chloride (136 mg, 1 mmol), 1*H*-1,2,3-triazole (83 mg, 1.2 mmol) and *N*,*N*-dimethyl-4-aminopyridine (DMAP) (cat.) in tetrahydrofuran (THF)/triethylamine (NEt_3_) (5:1, 2 mL) was stired overnight at 60 °C. The solvent was removed under vacuum and the resulting residue was purified on a silica gel column (Hexane/ethyl acetate (EtOAc) 7:3 to 6:4) giving KK002-N1 (**1**) as colorless oil (73mg, 43%) and KK002-N2 (**2**) as a white solid (55 mg, 33%).

### KK002-N1 (1)

^1^H NMR (500 MHz, CDCl_3_): δ 8.19 (d, *J* = 1.0 Hz, 1H), 7.72 (d, *J* = 1.0 Hz, 1H), 3.80-3.40 (m, 4H), 1.33 (t, *J* = 7.0 Hz, 6H). ^13^C NMR (125 MHz, CDCl_3_): δ 148.7, 132.7, 125.2, 44.3, 14.1. HRMS (*m*/*z*): calcd. for C_7_H_12_N_4_ONa [M+Na]^+^: 191.0903; found, 191.0905. **KK002-N2 (2)**; ^1^H NMR (500 MHz, CDCl_3_): δ 7.80 (s, 2H), 3.75-3.30 (m, 4H), 1.29 (t, *J* = 7.0 Hz, 6H). ^13^C NMR (125 MHz, CDCl_3_): δ 135.6, 43.1, 12.4. HRMS (*m/z*): calcd. for C_7_H_12_N_4_O [M+Na]^+^: 191.0903; found, 191.0911.

### KK003-N1 (3), KK003-N2 (4)

*N*,*N*-diphenyl-1*H*-1,2,3-triazole-1-carboxamide (**3**), *N*,*N-*diphenyl-2*H*-1,2,3-triazole-2-carboxamide (**4**).

The mixture of diphenylcarbamoyl chloride (280 mg, 1.21 mmol), 1*H*-1,2,3-triazole (100 mg, 1.45 mmol) and DMAP (cat.) in THF/NEt_3_ (5:1, 2 mL) was stired overnight at 60 °C. The solvent was removed under vacuum and the resulting residue was purified on a silica gel column (Hexane/EtOAc 7:3 to 6:4) giving KK003-N1 (**3**) (197 mg, 62%) as a white solid and KK003-N2 (**4**) (121mg, 38%) as a white solid.

### KK003-N1 (3)

^1^H NMR (500 MHz, CDCl_3_): δ 8.11 (d, *J* = 1.3 Hz, 1H), 7.59 (d, *J* = 1.3 Hz, 1H), 7.38-7.33 (m, 4H), 7.30-7.25 (m, 2H), 7.23-7.18 (m, 4H). ^13^C NMR (125 MHz, CDCl3): δ 148.81, 142.30, 132.74, 129.51, 127.49, 126.48, 124.51.

### KK003-N2 (4)

^1^H NMR (500 MHz, CDCl_3_): δ 8.11 (d, *J* = 1.3 Hz, 1H), 7.59 (d, *J* = 1.3 Hz, 1H), 7.38-7.33 (m, 4H), ^1^H NMR (500 MHz, CDCl_3_): δ 7.61 (s, 2H), 7.30-7.34 (m, 4H), 7.22-7.26 (m, 2H), 7.16-7.18 (m, 4H). ^13^C NMR (125 MHz, CDCl_3_): δ 149.31, 142.55, 136.17, 129.32, 127.02, 126.30.

### KK004-N1 (5), KK004-N2 (6)

morpholino(1*H*-1,2,3-triazol-1-yl)methanone (**5**), morpholino(2*H*-1,2,3-triazol-2-yl)methanone (**6**).

The mixture of 4-morpholinecarbonyl chloride (150 mg, 1 mmol), 1*H*-1,2,3-triazole (83 mg, 1.2 mmol) and DMAP (cat.) in THF/NEt_3_ (5:1, 2 mL) was stired overnight at 60 °C. The solvent was removed under vacuum and the resulting residue was purified on a silica gel column (Hexane/EtOAc 7:3 to 6:4) giving KK004-N1 (**5**) (53 mg, 29%) as a white solid and KK004-N2 (**6**) (26 mg, 14%) as a white solid.

### KK004-N1 (5)

^1^H NMR (500 MHz, CDCl_3_): δ 8.19 (d, *J* = 1.1 Hz, 1H), 7.74 (d, *J* = 1.1 Hz, 1H), 4.17-3.66 (m, 8H). ^13^C NMR (125 MHz, CDCl_3_): δ 133.0, 125.4, 66.6, 48.4, 45.7. HRMS (*m/z*): calcd. for C_7_H_10_N_4_O_2_ [M+Na]^+^: 205.0696; found, 205.0695.

### KK004-N2 (6)

1H NMR (500 MHz, CDCl_3_): δ 7.83 (s, 2H), 4.10-3.65 (m, 4H). ^13^C NMR (125 MHz, CDCl_3_): δ 136.2, 66.6, 48.4, 45.8. HRMS (*m/z*): calcd. for C_7_H_10_N_4_O_2_ [M+Na]^+^: 205.0696; found, 205.0694.

### KK007-N1 (7), KK007-N2 (8)

pyrrolidin-1-yl(1*H*-1,2,3-triazol-1-yl)methanone (**7**), pyrrolidin-1-yl(2*H*-1,2,3-triazol-2-yl)methanone (**8**).

The mixture of 1-pyrrolidinecarbonyl chloride (100 mg, 0.8 mmol), 1*H*-1,2,3-triazole (124 mg, 1.8 mmol) and DMAP (cat.) in THF/NEt_3_ (5:1, 2 mL) was stired overnight at 60 °C. The solvent was removed under vacuum and the resulting residue was purified on a silica gel column (Hexane/EtOAc 7:3 to 6:4) giving KK007-N1 (**7**) (68 mg, 55%) as a white solid and KK007-N2 (**8**) (36 mg, 29%) as a white solid.

### KK007-N1 (7)

^1^H NMR (500 MHz, CDCl_3_): δ 8.31 (d, *J* = 1.4 Hz, 1H), 7.72 (d, *J* = 1.4 Hz, 1H), 4.04 (t, *J* = 6.5 Hz, 2H), 3.74 (t, *J* = 6.7 Hz, 2H), 2.08-1.95 (m, 4H). ^13^C NMR (125 MHz, CDCl_3_): δ 147.12, 132.59, 124.65, 50.25, 48.92, 26.48, 23.90. HRMS (*m/z*): calcd. for C_7_H_10_N_4_O [M+Na]^+^: 189.0747; found, 189.0751.

### KK007-N2 (8)

^1^H NMR (500 MHz, CDCl_3_): δ 7.81 (s, 2H), 3.84 (t, *J* = 6.4 Hz, 2H), 3.75 (t, *J* = 6.5 Hz, 2H), 2.02-1.94 (m, 4H). ^13^C NMR (125 MHz, CDCl_3_): δ 135.80, 50.02, 48.62, 26.44, 24.12. HRMS (*m/z*): calcd. for C_7_H_10_N_4_O [M+Na]^+^: 189.0747; found, 189.0749.

### KK020 (9)

(4-benzyl-1*H*-1,2,3-triazol-1-yl)(pyrrolidin-1-yl)methanone.

The mixture of 1-pyrrolidinecarbonyl chloride (190 mg, 1.4 mmol), 4-benzyl-1*H*-1,2,3-triazole (226 mg, 1.4 mmol) and DMAP (cat.) in THF/NEt_3_ (5:1, 2 mL) was stired overnight at 60 °C. The solvent was removed under vacuum and the resulting residue was purified on a silica gel column (Hexane/EtOAc 7:3 to 6:4) giving KK020 (**9**) (86 mg, 24%) as yellow oil.

^1^H NMR (500 MHz, CDCl_3_): δ 7.91 (s, 1H), 7.21-7.34 (m, 5H), 4.11 (s, 2H), 4.01 (t, J = 6.3 Hz, 2H), 3.69 (t, *J* = 6.6 Hz, 2H), 1.92-2.04 (m, 4H). ^13^C NMR (125 MHz, CDCl_3_): δ 147.2, 146.5, 138.3, 128.7, 128.7, 126.7, 50.2, 48.9, 31.9, 26.5, 23.9. HRMS (*m/z*): calcd. for C_14_H_16_N_4_O [M+H]^+^: 257.1397; found, 257.1403.

### KK021 (10)

(4-(6-methoxynaphthalen-2-yl)-1*H*-1,2,3-triazol-1-yl)(pyrrolidin-1-yl)methanone. The mixture of 1-pyrrolidinecarbonyl chloride (190 mg, 1.4 mmol), 4-(6-methoxynaphthalen-2-yl)-1*H*-1,2,3-triazole (338 mg, 1.5 mmol) and DMAP (cat.) in THF/NEt_3_ (5:1, 2 mL) was stired overnight at 60 °C. The solvent was removed under vacuum and the resulting residue was purified on a silica gel column (Hexane/EtOAc 7:3 to 6:4) giving KK021 (**10**) (47 mg, 10%) as a white solid.

^1^H NMR (500 MHz, CDCl_3_): δ 8.56 (s, 1H), 8.34 (s, 1H), 7.92 (dd, *J* = 8.3, 1.7 Hz, 1H), 7.86-7.77 (m, 2H), 7.21-7.14 (m, 2H), 4.10 (t, *J* = 6.5 Hz, 2H), 3.94 (s, 3H), 3.77 (t, *J* = 6.5 Hz, 2H), 2.09-1.97 (m, 4H). ^13^C NMR (125 MHz, CDCl_3_): δ 158.13, 147.19, 146.62, 134.60, 129.81, 128.91, 127.51, 124.82, 124.79, 124.31, 120.01, 119.44, 105.78, 55.33, 50.34, 49.01, 26.58, 23.96. HRMS (*m/z*): calcd. for C_18_H_18_N_4_O_2_ [M+H]^+^: 323.1503; found, 323.1498.

### KK022 (11)

(4-phenyl-1*H*-1,2,3-triazol-1-yl)(pyrrolidin-1-yl)methanone.

The mixture of 1-pyrrolidinecarbonyl chloride (417 mg, 3.1 mmol), 4-phenyl-1*H*-1,2,3-triazole (300 mg, 2.1 mmol) and DMAP (cat.) in THF/NEt_3_ (5:1, 2 mL) was stired overnight at 60 °C. The solvent was removed under vacuum and the resulting residue was purified on a silica gel column (Hexane/ EtOAc 7:3 to 6:4) giving KK022 (**11**) (243 mg, 48%) as a white solid.

^1^H NMR (500 MHz, CDCl_3_): δ 8.49 (s, 1H), 7.91-7.86 (m, 2H), 7.49-7.43 (m, 2H), 7.41-7.34 (m, 1H), 4.07 (t, *J* = 6.7 Hz, 2H), 3.75 (t, *J* = 6.7 Hz, 2H), 2.09-1.95 (m, 4H). ^13^C NMR (125 MHz, CDCl_3_): δ147.08, 146.35, 129.60, 129.06, 128.91, 128.78, 128.60, 125.88, 120.17, 50.28, 48.96, 26.52, 23.89. HRMS (*m/z*): calcd. for C_13_H_14_N_4_O [M+H]^+^: 243.1240; found, 243.1238.

### KK025 (12)

(4-(4-phenoxyphenyl)-1*H*-1,2,3-triazol-1-yl)(pyrrolidin-1-yl)methanone.

The mixture of 1-pyrrolidinecarbonyl chloride (480 mg, 3.6 mmol), 4-(4-phenoxyphenyl)-1*H-*1,2,3-triazole (568 mg, 2.4 mmol) and DMAP (cat.) in THF/NEt_3_ (5:1, 2 mL) was stired overnight at 60 °C. The solvent was removed under vacuum and the resulting residue was purified on a silica gel column (Hexane/EtOAc 7:3 to 6:4) giving KK025 (**12**) (219 mg, 27%) as a white solid.

^1^H NMR (500 MHz, CDCl_3_): δ 8.44 (s, 1H), 7.84 (dt, *J* = 9.2, 2.4 Hz, 2H), 7.43-7.30 (m, 2H), 7.18-7.11 (m, 1H), 7.11-7.01 (m, 4H), 4.07 (t, *J* = 6.6 Hz, 2H), 3.75 (t, *J* = 6.6 Hz, 2H), 2.09-1.96 (m, 4H). ^13^C NMR (125 MHz, CDCl_3_): δ 157.80, 156.70, 147.09, 145.92, 129.82, 127.42, 124.61, 123.62, 119.68, 119.20, 118.98, 50.28, 48.97, 26.54, 23.90. HRMS (*m/z*): calcd. for C_13_H_12_N_4_OCl_2_ [M+H]^+^: 311.0461; found, 311.0460.

### KK030 (13)

(4-(3,4-dichlorophenyl)-1*H*-1,2,3-triazol-1-yl)(pyrrolidin-1-yl)methanone.

The mixture of 1-pyrrolidinecarbonyl chloride (810 mg, 6.1 mmol), 4-(3,4-dichlorophenyl)-1*H-*1,2,3-triazole(865mg, 4.0mmol) and DMAP (cat.) in THF/NEt_3_ (5:1, 2 mL) was stired overnight at 60 °C. The solvent was removed under vacuum and the resulting residue was purified on a silica gel column (Hexane/EtOAc 7:3 to 6:4) giving KK030 (**13**) (407 mg, 32%) as a white solid.

^1^H NMR (500 MHz, CDCl_3_): δ 8.51 (s, 1H), 8.00 (d, *J* = 1.9 Hz, 1H), 7.71 (dd, *J* = 8.3, 1.9 Hz, 1H), 7.52 (d, *J* = 8.3 Hz, 1H), 4.07 (t, *J* = 6.5 Hz, 2H), 3.76 (t, *J* = 6.7 Hz, 2H), 2.10-1.96 (m, 4H). ^13^C NMR (125 MHz, CDCl_3_): δ 146.76, 144.26, 133.20, 132.53, 130.94, 129.66, 127.68, 125.02, 120.75, 50.29, 49.06, 26.53, 23.90. HRMS (*m/z*): calcd. for C_19_H_18_N_4_O_2_ [M+H]^+^: 335.1503; found, 335.1505.

### KK031 (14)

pyrrolidin-1-yl(4-(4-(trifluoromethoxy)phenyl)-1*H*-1,2,3-triazol-1-yl)methanone. The mixture of 1-pyrrolidinecarbonyl chloride (432 mg, 3.2 mmol), 4-(4-(trifluoromethoxy)phenyl)-1*H*-1,2,3-triazole (494 mg, 2.2 mmol) and DMAP (cat.) in THF/NEt_3_ (5:1, 2 mL) was stired overnight at 60 °C. The solvent was removed under vacuum and the resulting residue was purified on a silica gel column (Hexane/EtOAc 7:3 to 6:4) giving KK031 (**14**) (264 mg, 38%) as a white solid.

^1^H NMR (500 MHz, CDCl_3_): δ 8.50 (s, 1H), 7.94-7.90 (m, 2H), 7.34-7.28 (m, 2H), 4.08 (t, *J* = 6.6 Hz, 2H), 3.76 (t, *J* = 6.6 Hz, 2H), 2.10-1.97 (m, 4H). ^13^C NMR (125 MHz, CDCl_3_): δ 149.29, 146.92, 145.14, 128.41, 127.34, 121.43, 120.42, 119.39, 50.30, 49.03, 26.54, 23.90. HRMS (*m/z*): calcd. for C_14_H_13_N_4_O_2_F_3_, [M+H]^+^: 327.1063, found, 327.1068.

### KK032 (15)

(4-(3,5-difluorophenyl)-1*H*-1,2,3-triazol-1-yl)(pyrrolidin-1-yl)methanone.

The mixture of 1-pyrrolidinecarbonyl chloride (817 mg, 6.1 mmol), 4-(3,5-difluorophenyl)-1*H-*1,2,3-triazole (739 mg, 4.1 mmol) and DMAP (cat.) in THF/NEt_3_ (5:1, 2 mL) was stired overnight at 60 °C. The solvent was removed under vacuum and the resulting residue was purified on a silica gel column (Hexane/EtOAc 7:3 to 6:4) giving KK032 (**15**) (264 mg, 38%) as a white solid.

^1^H NMR (500 MHz, CDCl_3_): δ 8.54 (s, 1H), 7.45-7.39 (m, 2H), 6.82 (tt, *J* = 8.8, 2.3 Hz, 1H), 4.07 (t, *J* = 6.6 Hz, 2H), 3.76 (t, *J* = 6.6 Hz, 2H), 2.10-1.97 (m, 4H). ^13^C NMR (125 MHz, CDCl_3_): δ 164.38, 162.40, 146.73, 144.45, 132.76, 121.15, 108.80, 103.84, 50.30, 49.07, 26.52, 23.89. HRMS (*m/z*): calcd. for C_13_H_12_N_4_OF_2_ [M+Na]^+^: 279.1052; found, 279.1054;

### KK052 (16)

The mixture of (4-phenylpiperazin-1-yl)(1*H*-1,2,3-triazol-1-yl)methanone (N1-isomer) and (4-phenylpiperazin-1-yl)(2*H*-1,2,3-triazol-2-yl)methanone (N2-isomer). The mixture of 4-phenylpiperazine-1-carbonyl chloride(191mg, 0.9mmol), 4-(3,5-difluorophenyl)-N*H*-1,2,3-triazole (70 mg, 1.0 mmol) and DMAP (cat.) in THF/NEt_3_ (5:1, 2 mL) was stired overnight at 60 °C. The solvent was removed under vacuum and the resulting residue was purified on a silica gel column (Hexane/EtOAc 7:3 to 6:4) giving KK052 (**16**) (215 mg, 98%, NMR ratio N1-isomer:N2-isomer=2:1) as a white solid.

### KK052-N1

^1^H NMR (500 MHz, CDCl_3_): δ 8.21 (d, *J* = 1.1 Hz, 1H), 7.75 (d, *J* = 1.1 Hz, 1H), 7.33-7.28 (m, 2H), 6.98-6.91 (m, 3H), 4.25-3.85 (m, 4H), 3.43-3.24 (m, 4H). ^13^C NMR (125 MHz, CDCl_3_): δ 150.58, 148.05, 132.99, 129.30, 125.44, 120.86, 116.75, 49.61, 49.31, 47.81, 45.47. HRMS (*m/z*): calcd. for C_13_H_15_N_5_O [M+H]^+^: 258.1349; found, 258.1347.

### KK052-N1

^1^H NMR (500 MHz, CDCl_3_) δ 7.85 (s, 2H), 7.33-7.28 (m, 2H), 6.97-6.91 (m, 3H), 4.03-3.83 (m, 4H), 3.44-3.19 (m, 4H). ^13^C NMR (126 MHz, CDCl_3_) δ 150.69, 148.93, 136.17, 129.28, 120.79, 116.76, 49.43, 47.56, 45.29. HRMS (*m/z*): calcd. for C_13_H_15_N_5_O [M+H]^+^: 258.1349; found, 258.1348.

### KK053-N1 (17)

(4-(2-chlorophenyl)piperazin-1-yl)(1*H*-1,2,3-triazol-1-yl)methanone. 1-(2-chlorophenyl)piperazine (150 mg, 0.8 mmol) and *N*,*N*-diisopropylethylamine (296 mg, 2.3 mmol) were added to the triphosgene (113 mg, 0.4 mmol) solution in THF (5 mL) keeping the tempareture inside <10 °C and cooled to 0 °C. The reaction was stirrer for 15 min on ice. After adding iced water, the reaction solution was extracted with EtOAc twice. The EtOAc layer was combined, dehydrated with Na_2_SO_4_ and the solvent was removed under vacuum and the carbamoyl chloride intermediate was obtained. The resulting residue was dissolved in THF (5 mL) and *N*,*N*-diisopropylethylamine (713 mg, 5.5 mmol), 1*H*-1,2,3-triazole (63 mg, 0.9 mmol) and DMAP (cat.) were added on ice and stirred overnight at room temperature (rt). Then the solvent was removed under vacuum and the resulting residue was dissolved and extracted with EtOAc and water twice. The EtOAc layer was combined and the solvent was removed under vacuum and the resulting residue was purified on a silica gel column (Hexane:AcOEt=7:3 to 1:1) giving KK053-N1 (**17**) (88 mg, 40%) as a white solid. ^1^H NMR (500 MHz, CDCl_3_): δ 8.21 (d, *J* = 1.3 Hz, 1H), 7.75 (d, *J* = 1.3 Hz, 1H), 7.40 (dd, *J* = 7.8, 1.4 Hz, 1H), 7.27-7.23 (m, 1H), 7.08-7.01 (m, 2H), 4.27-3.86 (m, 4H), 3.28-3.13 (m, 4H). ^13^C NMR (125 MHz, CDCl_3_): δ 148.24, 132.97, 130.75, 128.97, 127.74, 125.40, 124.54, 120.61, 51.26, 50.94, 48.22, 45.82. HRMS (*m/z*): calcd. for C_13_H_14_N_5_OCl [M+H]^+^: 292.0960; found, 292.0958.

### KK054-N1 (18)

(4-(4-chlorophenyl)piperazin-1-yl)(1*H*-1,2,3-triazol-1-yl)methanone. KK054-N1 (**18**) was synthesized as the same method as compound **17**. The carbamoyl chloride intermediate was obtained with 1-(4-chlorophenyl)piperazine (150 mg, 0.8 mmol), *N*,*N-*diisopropylethylamine (296 mg, 2.3 mmol), triphosgene (113 mg, 0.4 mmol) and THF (5 mL). Then KK054-N1 (**18**) (86 mg, 39%) was obtained as a white solid with the carbamoyl chloride intermediate, THF (5 mL), *N*,*N*-diisopropylethylamine (713 mg, 5.5 mmol), 1*H*-1,2,3-triazole (63 mg, 0.9 mmol) and DMAP (cat.). ^1^H NMR (500 MHz, CDCl_3_): δ 8.21 (d, *J* = 1.3 Hz, 1H), 7.75 (d, *J* = 1.3 Hz, 1H), 7.26-7.22 (m, 2H), 6.89-6.85 (m, 2H), 4.26-3.85 (m, 4H), 3.41-3.19 (m, 4H). ^13^C NMR (125 MHz, CDCl_3_): δ 149.21, 148.00, 133.00, 129.16, 125.76, 125.43, 117.93, 49.58, 49.28, 47.67, 45.33. HRMS (*m/z*): calcd. for C_13_H_14_N_5_OCl [M+H]^+^: 292.0960; found, 292.0960.

### KK055-N1 (19)

(4-(pyridin-2-yl)piperazin-1-yl)(1*H*-1,2,3-triazol-1-yl)methanone.

KK055-N1 (**19**) was synthesized as the same method as compound **17**. The carbamoyl chloride intermediate was obtained with 1-(2-pyridyl)piperazine (150 mg, 0.9 mmol), *N*,*N-*diisopropylethylamine (356 mg, 2.8 mmol), triphosgene (136 mg, 0.5 mmol) and THF (5 mL). Then KK055-N1 (**19**) (61 mg, 26%) was obtained as a white solid with the carbamoyl chloride intermediate, THF (5 mL), *N*,*N*-diisopropylethylamine (356 mg, 2.8 mmol), 1*H*-1,2,3-triazole (76 mg, 1.1 mmol) and DMAP (cat.). 1H NMR (500 MHz, CDCl_3_): δ 8.27-8.15 (m, 2H), 7.75 (d, *J* = 1.1 Hz, 1H), 7.57-7.51 (m, 1H), 6.73-6.65 (m, 2H), 4.25-3.83 (m, 4H), 3.83-3.61 (m, 4H). ^13^C NMR (125 MHz, CDCl_3_): δ 158.79, 148.19, 148.03, 137.78, 133.00, 125.42, 114.21, 107.21, 47.53, 45.27, 44.73. HRMS (*m/z*): calcd. for C_12_H_14_N_6_O [M+H]^+^: 259.1302; found, 259.1304.

### KK067 (20)

The mixture of (4-(4-nitrophenyl)piperazin-1-yl)(1*H*-1,2,3-triazol-1-yl)methanone (N1 isomer) and (4-(4-nitrophenyl)piperazin-1-yl)(2*H*-1,2,3-triazol-2-yl)methanone (N1 isomer). KK067 (**20**) was synthesized as the same method as compound **17**. The carbamoyl chloride intermediate was obtained with 1-(4-nitrophenyl)piperazine (100 mg, 0.5 mmol), *N*,*N-*diisopropylethylamine (187 mg, 1.5 mmol), triphosgene (72 mg, 0.2 mmol) and THF (5 mL).

Then the KK067-N1, N2 mixture (**20**) (97 mg, 66%; NMR ratio N1-isomer:N2-isomer = 6:4) was obtained as a yellow solid with the carbamoyl chloride intermediate, THF (5 mL), *N*,*N-*diisopropylethylamine (187 mg, 1.5 mmol), 1*H*-1,2,3-triazole (40 mg, 0.6 mmol) and DMAP (cat.).

### KK067-N1

1H NMR (500 MHz, CDCl_3_): δ 8.24 (d, *J* = 1.3 Hz, 1H), 8.22-8.12 (m, 2H), 7.77 (d, *J* = 1.3 Hz, 1H), 6.92-6.82 (m, 2H), 4.32-3.91 (m, 4H), 3.69-3.53 (m, 4H).

### KK067-N2

^1^H-NMR (500 MHz, CDCl_3_): δ 8.22-8.12 (m, 2H), 7.87 (s, 2H), 6.92-6.82 (m, 2H), 4.32-3.91 (m, 4H), 3.69-3.53 (m, 4H). ^13^C NMR (125 MHz, CDCl_3_): δ 154.26, 154.19, 148.86, 148.06, 139.40, 139.32, 136.46, 133.12, 125.94, 125.49, 113.15, 113.12, 106.46, 47.01. HRMS (*m/z*): calcd. for C_13_H_14_N_6_O_3_ [M+H]^+^: 303.1200; found, 303.1201.

### KK070 (21)

The mixture of 4-(4-(1*H*-1,2,3-triazole-1-carbonyl)piperazin-1-yl)benzonitrile (N1-isomer) and 4-(4-(2*H*-1,2,3-triazole-2-carbonyl)piperazin-1-yl)benzonitrile (N2-isomer). KK070 (**21**) was synthesized as the same method as compound **17**. The carbamoyl chloride intermediate was obtained with 4-(1-piperazinyl)benzonitrile (150 mg, 0.8 mmol), *N*,*N-*diisopropylethylamine (311 mg, 2.4 mmol), triphosgene (119 mg, 0.4 mmol) and THF (5 mL). Then the KK070-N1, N2 mixture (**21**) (117mg, 52%; NMR ratio N1-isomer:N2-isomer = 87:12) was obtained as a white solid with the carbamoyl chloride intermediate, THF (5 mL), *N*,*N-*diisopropylethylamine (311 mg, 2.4 mmol), 1*H*-1,2,3-triazole (66 mg, 1.0 mmol) and DMAP (cat.).

### KK070-N1

^1^H NMR (500 MHz, CDCl_3_): δ 8.23 (d, *J* = 1.3 Hz, 1H), 7.76 (d, *J* = 1.3 Hz, 1H), 7.59-7.51 (2H, m), 6.94-6.87 (m, 2H), 4.30-3.88 (m, 4H), 3.59-3.43 (m, 4H).

### KK070-N2

^1^H-NMR (500 MHz, CDCl_3_): δ 7.86 (s, 2H), 7.59-7.51 (m, 2H), 6.94-6.87 (m, 2H), 4.30-3.88 (m, 4H), 3.59-3.43 (m, 4H). ^13^C NMR (125 MHz, CDCl_3_): δ 152.69, 148.00, 136.37, 133.60, 133.06, 125.45, 119.57, 114.64, 101.61, 47.22, 44.88. HRMS (*m/z*): calcd. for C_14_H_14_N_6_O [M+H]^+^: 283.1302; found, 283.1303.

### KK071 (22)

The mixture of (4-cyclohexylpiperazin-1-yl)(1*H*-1,2,3-triazol-1-yl)methanone (N1-isomer) and (4-cyclohexylpiperazin-1-yl)(2*H*-1,2,3-triazol-2-yl)methanone (N2-isomer). KK071 (**22**) was synthesized as the same method as compound **17**. The carbamoyl chloride intermediate was obtained with 1-cyclohexylpiperazine (150 mg, 0.9 mmol), *N*,*N-*diisopropylethylamine (346 mg, 2.7 mmol), triphosgene (132 mg, 0.5 mmol) and THF (5 mL). Then the KK071-N1, N2 mixture (**22**) (169 mg, 72%, NMR ratio N1-isomer:N2-isomer = 64:36) was obtained as colorless oil with the carbamoyl chloride intermediate, THF (5 mL), *N*,*N-* diisopropylethylamine (346 mg, 2.7 mmol), 1*H*-1,2,3-triazole (74 mg, 1.1 mmol) and DMAP (cat.).

### KK071-N1

^1^H-NMR (500 MHz, CDCl_3_): δ 8.17 (d, *J* = 1.3 Hz, 1H), 7.73 (d, *J* = 1.3 Hz, 1H), 4.01-3.61 (m, 4H), 2.81-2.59 (m, 4H), 2.43-2.25 (m, 2H), 1.93-1.75 (m, 4H), 1.70-1.56 (m, 1H), 1.36-0.99 (m, 4H).

### KK071-N2

^1^H-NMR (500 MHz, CDCl_3_): δ 7.82 (s, 2H), 4.01-3.61 (m, 4H), 2.81-2.59 (m, 4H), 2.43-2.25 (m, 2H), 1.93-1.75 (m, 4H), 1.70-1.56 (m, 1H), 1.36-0.99 (m, 4H). ^13^C NMR (125 MHz, CDCl_3_): δ 148.88, 147.96, 135.87, 132.84, 125.27, 63.46, 49.06, 48.39, 45.97, 28.81, 26.16, 25.73. HRMS (*m/z*): calcd. for C_13_H_21_N_5_O [M+H]^+^: 264.1819; found, 264.1830.

### KK072 (23)

The mixture of (4-(p-tolyl)piperazin-1-yl)(1*H*-1,2,3-triazol-1-yl)methanone (N1-isomer) and (4-(p-tolyl)piperazin-1-yl)(2*H*-1,2,3-triazol-2-yl)methanone (N2-isomer)

KK072 (**23**) was synthesized as the same method as compound **17**. The carbamoyl chloride intermediate was obtained with 1-(4-methylphenyl)piperazine (150 mg, 0.9 mmol), *N*,*N-*diisopropylethylamine (330 mg, 2.6 mmol), triphosgene (71 mg, 1.0 mmol) and THF (5 mL). Then the KK072-N1, N2 mixture (**23**) (231 mg, quant, NMR ratio N1-isomer:N2-isomer = 69:31) was obtained as a white solid with the carbamoyl chloride intermediate, THF (5 mL), *N*,*N-*diisopropylethylamine (330 mg, 2.6 mmol), 1*H*-1,2,3-triazole (71 mg, 1.0 mmol) and DMAP (cat.).

### KK072-N1

^1^H NMR (500 MHz, CDCl_3_): δ 8.20 (d, *J* = 1.1 Hz, 1H), 7.74 (d, *J* = 1.1 Hz, 1H), 7.13-7.09 (m, 2H), 6.90-6.84 (m, 2H), 4.25-3.80 (m, 4H), 3.45-3.10 (m, 4H), 2.29 (s, 3H). **KK072-N2**: ^1^H NMR (500 MHz, CDCl_3_): δ 7.84 (s, 1H), 7.13-7.09 (m, 2H), 6.90-6.84 (m, 2H), 4.25-3.80 (m, 4H), 3.45-3.10 (m, 4H), 2.29 (s, 3H). ^13^C NMR (125 MHz, CDCl_3_): δ 148.57, 148.47, 148.04, 136.11, 132.95, 130.48, 130.42, 129.80, 125.41, 117.09, 49.92, 47.74, 45.47, 20.42. HRMS (*m/z*): calcd. for C_14_H_17_N_5_O [M+H]^+^: 272.1506; found, 272.1517.

### KK073 (24)

The mixture of (1*H*-1,2,3-triazol-1-yl)(4-(4-(trifluoromethyl)phenyl)piperazin-1-yl)methanone (N1-isomer) and (2*H*-1,2,3-triazol-2-yl)(4-(4-(trifluoromethyl)phenyl)piperazin-1-yl)methanone (N2-isomer).

KK073 (**24**) was synthesized as the same method as compound **17**. The carbamoyl chloride intermediate was obtained with 1-(4-trifluoromethylphenyl)piperazine (150 mg, 0.7 mmol), *N*,*N-*diisopropylethylamine (253 mg, 2.0 mmol), triphosgene (97 mg, 0.3 mmol) and THF (5 mL). Then the KK073-N1, N2 mixture (**24**) (188 mg, 89%, NMR ratio N1-isomer:N2-isomer = 73:17) was obtained as a white solid with the carbamoyl chloride intermediate, THF (5 mL), *N*,*N-* diisopropylethylamine (253 mg, 2.0 mmol), 1*H*-1,2,3-triazole (54 mg, 0.8 mmol) and DMAP (cat.).

### KK073-N1

^1^H NMR (500 MHz, CDCl_3_): δ 8.22 (d, *J* = 1.4 Hz, 1H), 7.76 (d, *J* = 1.4 Hz, 1H), 7.56-7.50 (m, 2H), 6.98-6.94 (m, 2H), 3.95-4.18 (m, 4H), 3.45-3.45 (m, 4H).

### KK073-N2

^1^H NMR (500 MHz, CDCl_3_): δ 7.86 (s, 1H), 7.56-7.50 (m, 2H), 6.98-6.94 (m, 2H), 4.18-3.95 (m, 4H), 3.56-3.32 (m, 4H). ^13^C NMR (125 MHz, CDCl_3_): δ 152.72, 152.64, 148.90, 148.06, 136.31, 133.07, 127.74, 126.59, 125.59, 125.48, 123.44, 121.87, 115.18, 77.28, 77.03, 76.77, 48.31, 47.49, 45.12. HRMS (*m/z*): calcd. for C_14_H_14_N_5_OF_3_ [M+H]^+^: 326.1223; found, 326.1233.

### KK075 (25)

The mixture of (4-benzhydrylpiperazin-1-yl)(1*H*-1,2,3-triazol-1-yl)methanone (N1-isomer) and (4-benzhydrylpiperazin-1-yl)(2*H*-1,2,3-triazol-2-yl)methanone (N2-isomer). KK075 (**25**) was synthesized as the same method as compound **17**. The carbamoyl chloride intermediate was obtained with 1-(diphenylmethyl)piperazine (150 mg, 0.6 mmol), *N*,*N-*diisopropylethylamine (230 mg, 1.8 mmol), triphosgene (97 mg, 0.3 mmol) and THF (5 mL). Then the KK075-N1, N2 mixture (**25**) (186 mg, 90%, NMR ratio N1-isomer:N2-isomer = 65:35) was obtained as a white solid with the carbamoyl chloride intermediate, THF (5 mL), *N*,*N-*diisopropylethylamine (230 mg, 1.8 mmol), 1*H*-1,2,3-triazole (49 mg, 0.7 mmol) and DMAP (cat.).

### KK075-N1

^1^H NMR (500 MHz, CDCl_3_): δ 8.13 (d, *J* = 1.1 Hz, 1H), 7.69 (d, *J* = 1.1 Hz, 1H), 7.45-7.39 (m, 4H), 7.32-7.25 (m, 4H), 7.23-7.17 (2H), 4.31 (s, 1H), 4.07-3.55 (m, 4H), 2.69-2.36 (m, 4H).

### KK075-N2

^1^H NMR (500 MHz, CDCl_3_): δ 7.77 (s, 2H), 7.45-7.39 (m, 4H), 7.32-7.25 (m, 4H), 7.23-7.17 (2H), 4.29 (s, 1H), 4.07-3.55 (m, 4H), 2.69-2.36 (m, 4H). ^1^3C NMR (125 MHz, CDCl_3_): δ 148.91, 148.01, 141.91, 141.76, 135.86, 132.84, 128.65, 127.83, 127.26, 127.22, 125.24, 75.84, 75.77, 51.83, 51.28, 47.92, 45.48. HRMS (*m/z*): calcd. for C_20_H_21_N_5_O [M+H]^+^: 348.1819; found, 348.1834.

### KK094 (26)

The mixture of indolin-1-yl(1*H*-1,2,3-triazol-1-yl)methanone (N1-isomer) and indolin-1-yl(2*H*-1,2,3-triazol-2-yl)methanone (N2-isomer).

KK094 (**26**) was synthesized as the same method as compound **17**. The carbamoyl chloride intermediate was obtained with indoline (400 mg, 3.4 mmol), *N*,*N*-diisopropylethylamine (1.301 g, 10.1 mmol), triphosgene (498 mg, 1.7 mmol) and THF (5 mL). Then the KK094-N1, N2 mixture (**26**) (675 mg, 94%, NMR ratio N1-isomer:N2-isomer = 6:4)was obtained as a white solid with the carbamoyl chloride intermediate, THF (10 mL), *N*,*N*-diisopropylethylamine (1.301 g, 10.1 mmol), 1*H*-1,2,3-triazole (49 mg, 0.7 mmol) and DMAP (cat.). Then the mixture was further purified on a silica gel column (Hexane:CH_2_Cl_2_:EtOAc = 1:4:0.4 to 0:4:0.4) giving KK094-N1 (190 mg, 26%) as a white solid and KK094-N2 (97 mg, 14%) as a white solid.

### KK094-N1

^1^H NMR (500 MHz, CDCl_3_): δ 8.36 (d, *J* = 1.1 Hz, 1H), 8.11 (br s, 1H), 7.78 (d, *J* = 1.1 Hz, 1H), 7.33-7.27 (m, 2H), 7.16 (t, *J* = 7.4 Hz, 1H), 4.67 (t, J = 8.3 Hz, 2H), 3.26 (t, J = 8.3 Hz, 2H). ^13^C NMR (125 MHz, CDCl_3_): δ 145.92, 141.88, 132.91, 132.19, 127.68, 125.45, 125.00, 124.96, 117.49, 51.77, 28.55. HRMS (*m*/*z*): calcd. for C11H10N4O, [M+H]^+^: 215.0927; found, 215.0927.

### KK094-N2

^1^H NMR (500 MHz, CDCl_3_): δ 8.10 (s, 1H), 7.88 (s, 2H), 7.35-7.20 (m, 2H), 7.13 (t, *J* = 7.4 Hz, 1H), 4.47 (t, *J* = 8.3 Hz, 2H), 3.22 (t, *J* = 8.3 Hz, 2H). ^13^C NMR (125 MHz, CDCl_3_): δ 146.35, 141.95, 136.27, 132.03, 127.67, 125.04, 124.84, 117.48, 51.39, 28.45. HRMS (m/z): calcd. for C11H10N4O [M+Na]^+^:237.0747; found,237.0757.

### KK099 (27)

The mixture of isoindolin-2-yl(1*H*-1,2,3-triazol-1-yl)methanone (N1-isomer) and isoindolin-2-yl(2*H*-1,2,3-triazol-2-yl)methanone (N2-isomer).

KK099 (**27**) was synthesized as the same method as compound **17**. The carbamoyl chloride intermediate was obtained with isoindoline (200 mg, 1.7 mmol), *N*,*N*-diisopropylethylamine (651 mg, 5.0 mmol), triphosgene (249 mg, 0.9 mmol) and THF (5 mL). Then the KK099-N1, N2 mixture (**27**) (675 mg, 94%, NMR ratio N1-isomer:N2-isomer = 6:4) was obtained as a white solid with the carbamoyl chloride intermediate, THF (10 mL), *N*,*N*-diisopropylethylamine (651 mg, 5.0 mmol), 1*H*-1,2,3-triazole (116 mg, 2.0 mmol) and DMAP (cat.). Then the mixture was further purified on a silica gel column (Hexane:CH_2_Cl_2_:EtOAc = 1:4:0.4 to 0:4:0.4) giving KK099-N1 (93 mg, 26%) as a white solid an099-N2 (86 mg, 24%) as a white solid.

### KK099-N1

^1^H NMR (500 MHz, CDCl_3_): δ 8.36 (d, *J* = 1.1 Hz, 1H), 8.11 (br s, 1H), 7.78 (d, *J* = 1.1 Hz, 1H), 7.33-7.27 (m, 2H), 7.16 (t, *J* = 7.4 Hz, 1H), 4.67 (t, J = 8.3 Hz, 2H), 3.26 (t, J = 8.3 Hz, 2H). ^1^H NMR (500 MHz, CDCl_3_): δ 8.40 (d, *J* = 1.3 Hz, 1H), 7.77 (d, *J* = 1.3 Hz, 1H), 7.38-7.28 (m, 4H), 5.47 (s, 2H), 5.11 (s, 2H). ^13^C NMR (125 MHz, CDCl_3_): δ 147.1, 136.4, 134.0, 132.8, 128.0, 127.9, 124.9, 122.7, 122.6, 56.0, 55.1. HRMS (*m/z*): calcd. for C_11_H_10_N_4_O, [M+H]^+^: 215.0927; found, 215.0920.

### KK099-N2

^1^H NMR (500 MHz, CDCl_3_): δ 7.89 (s, 2H), 7.37-7.24 (m, 5H), 5.31 (s, 2H), 5.13 (s, 2H). ^13^C NMR (125 MHz, CDCl_3_): δ 147.7, 136.4, 136.3, 134.6, 127.9, 127.8, 122.6, 122.5, 55.8, 55.0. HRMS (*m/z*): calcd. for C_11_H_10_N_4_O [M+Na]^+^: 237.0747; found, 237.0749.

### KK122 (28)

(1*H*-imidazol-1-yl)(indolin-1-yl)methanone. 1,1’-carbonyldiimidazole (180 mg, 1.1 mmol) was added to the solution of indoline (100 mg, 0.8 mmol) in THF (5 mL) and strred overnight at rt. Then the solvent was removed under vacuum and the resulting residue was dissolved in EtOAc and water and pH wasadjusted to 7 with 0.5 M HCl, and extracted with EtOAc and water twice. The EtOAc layer was combined and the solvent was removed under vacuum and the resulting residue was purified on a silica gel column (EtOAc:MeOH = 1:0 to 9:1) giving KK122 (**28**) (147 mg, 82%) as a white solid.

^1^H NMR (500 MHz, CDCl_3_): δ 8.02 (m, 1H), 7.41 (br s, 1H), 7.36 (m, 1H), 7.26 (d, *J* = 7.5 Hz, 1H), 7.20 (t, *J* = 7.5 Hz, 1H), 7.13-7.13 (m, 1H), 7.09 (td, *J* = 7.5, 0.8 Hz, 1H), 4.19 (t, *J* = 8.2 Hz, 2H), 3.20 (t, *J* = 8.2 Hz, 2H). ^13^C NMR (125 MHz, CDCl_3_): δ 148.16, 141.30, 136.58, 131.91, 129.81, 127.56, 125.07, 124.87, 117.45, 116.44, 50.97, 28.30.

### Compound 29

(2-methylindolin-1-yl)(1*H*-1,2,3-triazol-1-yl)methanone.

Compound **29** was synthesized as the same method as compound **17**. The carbamoyl chloride intermediate was obtained with 2-methylindoline (300 mg, 2.3 mmol), *N*,*N*-diisopropylethylamine (2.08 g, 16.1 mmol2.08g, 16.1mmol), triphosgene (267 mg, 0.9 mmol) and THF (5 mL). Then the compound **29** (55 mg, 11%) was obtained as colorless oil with the carbamoyl chloride intermediate, THF (5 mL), *N*,*N*-diisopropylethylamine (873 mg, 6.8 mmol), 1*H*-1,2,3-triazole (187 mg, 2.7 mmol) and DMAP (cat.). ^1^H NMR (500 MHz, CDCl_3_): δ 8.35 (d, *J* = 1.1 Hz, 1H), 8.09 (br s, 1H), 7.78 (d, *J* = 1.1 Hz, 1H), 7.38-7.27 (m, 2H), 7.17 (t, *J* = 7.2 Hz, 1H), 5.70 (m, 1H), 3.53 (dd, *J* = 15.5, 9.2 Hz, 1H), 2.78 (d, *J* = 15.5 Hz, 1H), 1.27 (d, *J* = 6.3 Hz, 3H). ^13^C NMR (125 MHz, CDCl_3_): δ 145.76, 140.88, 132.87, 131.29, 127.69, 125.54, 125.67, 125.13, 118.00, 58.43, 36.47, 21.56. HRMS (*m/z*): calcd. for C12H12N4O [M+Na]^+^: 251.0903; found, 251.904.

### Compound 30

(3-methylindolin-1-yl)(1*H*-1,2,3-triazol-1-yl)methanone.

Compound **30** was synthesized as the same method as compound **17**. The carbamoyl chloride intermediate was obtained with 3-methylindoline (713 mg, 5.4 mmol), *N*,*N*-diisopropylethylamine (2.08 g, 16.1 mmol), triphosgene (635 mg, 2.1 mmol) and THF (5 mL). Then the compound **30** (509 mg, 42%) was obtained as a white solid with the carbamoyl chloride intermediate, THF (5 mL), *N*,*N*-diisopropylethylamine (2.08 g, 16.1 mmol), 1*H*-1,2,3-triazole (187 mg, 2.7 mmol) and DMAP (cat.). ^1^H NMR (500 MHz, CDCl_3_): δ 8.37 (d, *J* = 1.1 Hz, 1H), 8.09 (br s, 1H), 7.78 (d, *J* = 1.1 Hz, 1H), 7.35-7.22 (m, 2H), 7.19 (t, *J* = 7.4 Hz, 1H), 4.82 (dd, *J* = 11.5, 9.7 Hz, 1H), 4.22 (dd, *J* = 11.5, 6.9 Hz, 1H), 3.62-3.51 (m, 1H), 1.39 (d, *J* = 6.9 Hz, 3H). ^13^C NMR (125 MHz, CDCl_3_): δ 145.81, 141.43, 137.37, 132.92, 127.85, 125.60, 124.97, 123.81, 117.38, 59.55, 35.28, 19.64. HRMS (*m/z*): calcd. for C12H12N4O [M+H]^+^: 229.1084; found, 229.1085.

### Compound 31

(4-nitroindolin-1-yl)(1*H*-1,2,3-triazol-1-yl)methanone.

Compound **31** was synthesized as the same method as compound **17**. The carbamoyl chloride intermediate was obtained with 4-nitroindoline (837 mg, 5.1 mmol), *N*,*N*-diisopropylethylamine (1.98 g, 15.3 mmol), triphosgene (605 mg, 2.0 mmol) and THF (5 mL). Then the compound **31** (64 mg, 5%) was obtained as a yellow solid with the carbamoyl chloride intermediate, THF (5 mL), *N*,*N*-diisopropylethylamine (1.98 g, 15.3 mmol), 1*H*-1,2,3-triazole (423 mg, 6.1 mmol) and DMAP (cat.). ^1^H NMR (500 MHz, CDCl_3_): δ 8.45 (br s, 1H), 8.39 (d, *J* = 1.1 Hz, 1H), 8.02 (d, *J* = 8.3 Hz, 1H), 7.81 (d, *J* = 1.1 Hz, 1H), 7.51 (t, *J* = 8.3 Hz, 1H), 4.80 (t, *J* = 8.4 Hz, 2H), 3.77 (t, *J* = 8.4 Hz, 2H). ^13^C NMR (125 MHz, CDCl_3_): δ 146.34, 145.26, 144.67, 133.19, 129.57, 129.12, 125.10, 122.82, 120.54, 52.08, 29.78. HRMS (*m/z*): calcd. for C11H9N5O3 [M+H]^+^: 260.0778; found, 260.0781.

### Compound 32

(4-chloroindolin-1-yl)(1*H*-1,2,3-triazol-1-yl)methanone.

Compound **32** was synthesized as the same method as compound **17**. The carbamoyl chloride intermediate was obtained with 4-chloroindoline (1.00 g, 6.5 mmol), *N*,*N*-diisopropylethylamine (2.52 g, 19.5 mmol), triphosgene (773 mg, 2.6 mmol) and THF (5 mL). Then the compound **32** (47 mg, 3%) was obtained as a white solid with the carbamoyl chloride intermediate, THF (5 mL), *N*,*N*-diisopropylethylamine (2.52 g, 19.5 mmol), 1*H*-1,2,3-triazole (540 mg, 7.8 mmol) and DMAP (cat.). 1H NMR (500 MHz, CDCl_3_): δ 8.36 (d, *J* = 1.1 Hz, 1H), 8.01 (br s, 1H), 7.78 (d, *J* = 1.1 Hz, 1H), 7.28-7.22 (m, 2H), 7.15 (d, *J* = 7.4 Hz, 1H), 4.72 (t, *J* = 8.3 Hz, 2H), 3.28 (t, *J* = 8.3 Hz, 2H). ^13^C NMR (125 MHz, CDCl_3_): δ 146.03, 143.22, 136.51, 133.00, 130.88, 129.18, 125.34, 125.02, 115.72, 51.63, 28.05. HRMS (*m/z*): calcd. for C11H9N4OCl [M+H]^+^: 249.0538; found, 249.0538.

### Compound 33

(4-methylindolin-1-yl)(1*H*-1,2,3-triazol-1-yl)methanone.

Compound **33** was synthesized as the same method as compound **17**. The carbamoyl chloride intermediate was obtained with 4-methylindoline (730 mg, 5.5 mmol), *N*,*N*-diisopropylethylamine (2.13 g, 16.4 mmol), triphosgene (651 mg, 2.2 mmol) and THF (5 mL). Then the compound **33** (423 mg, 34%) was obtained as a white solid with the carbamoyl chloride intermediate, THF (5 mL), *N*,*N*-diisopropylethylamine (2.13g, 16.4mmol), 1*H*-1,2,3-triazole (454 mg, 6.6 mmol) and DMAP (cat.). 1H NMR (500 MHz, CDCl_3_): δ 8.36 (d, *J* = 1.1 Hz, 1H), 7.95 (br s, 1H), 7.77 (d, *J* = 1.1 Hz, 1H), 7.21 (t, *J* = 7.7 Hz, 1H), 6.99 (d, *J* = 7.4 Hz, 1H), 4.68 (t, *J* = 8.2 Hz, 2H), 3.15 (t, *J* = 8.2 Hz, 2H), 2.29 (s, 3H). ^13^C NMR (125 MHz, CDCl_3_): δ 145.91, 141.60, 134.48, 132.87, 130.96, 127.81, 126.40, 124.95, 114.90, 51.68, 27.46, 18.62. HRMS (*m/z*): calcd. for C12H12N4O, [M+Na]^+^: 251.0903; found, 251.0907.

### Compound 34

(5-nitroindolin-1-yl)(1*H*-1,2,3-triazol-1-yl)methanone.

Compound **34** was synthesized as the same method as compound **17**. The carbamoyl chloride intermediate was obtained with 5-nitroindoline (1.00 g, 6.1 mmol), *N*,*N*-diisopropylethylamine (2.36 g, 18.3 mmol), triphosgene (723 mg, 2.4 mmol) and THF (5 mL). Then the compound **34** (143 mg, 9%) was obtained as a yellow solid with the carbamoyl chloride intermediate, THF (5 mL), *N*,*N*-diisopropylethylamine (2.36 g, 18.3 mmol), 1*H*-1,2,3-triazole (505 mg, 7.3 mmol) and DMAP (cat.). 1H NMR (500 MHz, CDCl_3_): δ 8.40 (d, *J* = 1.3 Hz, 1H), 8.26-8.19 (m, 2H), 8.16 (t, *J* = 1.4 Hz, 1H), 7.81 (d, *J* = 1.1 Hz, 1H), 4.84 (t, *J* = 8.5 Hz, 2H), 3.38 (t, *J* = 8.5 Hz, 2H). ^13^C NMR (125 MHz, CDCl_3_): δ 147.65, 146.31, 145.04, 133.61, 133.26, 125.22, 124.44, 120.61, 117.25, 52.64, 28.12. HRMS (*m/z*): calcd. for C11H9N5O3 [M+Na]^+^: 282.0598; found, 282.0596.

### Compound 35

(5-chloroindolin-1-yl)(1*H*-1,2,3-triazol-1-yl)methanone.

Compound **35** was synthesized as the same method as compound **17**. The carbamoyl chloride intermediate was obtained with 5-chloroindoline (1.00 g, 6.5 mmol), *N*,*N*-diisopropylethylamine (2.52 g, 19.5 mmol), triphosgene (773 mg, 2.6 mmol) and THF (5 mL). Then the compound **35** (450 mg, 28%) was obtained as a white solid with the carbamoyl chloride intermediate, THF (5 mL), *N*,*N*-diisopropylethylamine (2.52 g, 19.5 mmol), 1*H*-1,2,3-triazole (540 mg, 7.8 mmol) and DMAP (cat.). 1H NMR (500 MHz, CDCl_3_): δ 8.36 (d, *J* = 1.1 Hz, 1H), 8.04 (br s, 1H), 7.77 (d, *J* = 1.1 Hz, 1H), 4.70 (t, *J* = 8.3 Hz, 2H), 3.25 (t, *J* = 8.3 Hz, 2H). ^13^C NMR (125 MHz, CDCl_3_): δ 145.88, 140.65, 134.03, 132.98, 130.51, 127.73, 125.17, 124.99, 118.39, 51.95, 28.37. HRMS (*m/z*): calcd. for C11H9N4OCl [M+H]^+^: 249.0538; found, 249.0538.

### Compound 35

(5-chloroindolin-1-yl)(1*H*-1,2,3-triazol-1-yl)methanone.

Compound **35** was synthesized as the same method as compound **17**. The carbamoyl chloride intermediate was obtained with 5-chloroindoline (1.00 g, 6.5 mmol), *N*,*N*-diisopropylethylamine (2.52 g, 19.5 mmol), triphosgene (773 mg, 2.6 mmol) and THF (5 mL). Then the compound **35** (450 mg, 28%) was obtained as a white solid with the carbamoyl chloride intermediate, THF (5 mL), *N*,*N*-diisopropylethylamine (2.52 g, 19.5 mmol), 1*H*-1,2,3-triazole (540 mg, 7.8 mmol) and DMAP (cat.). 1H NMR (500 MHz, CDCl_3_): δ 8.36 (d, *J* = 1.1 Hz, 1H), 8.04 (br s, 1H), 7.77 (d, *J* = 1.1 Hz, 1H), 4.70 (t, *J* = 8.3 Hz, 2H), 3.25 (t, *J* = 8.3 Hz, 2H). ^13^C NMR (125 MHz, CDCl_3_): δ 145.88, 140.65, 134.03, 132.98, 130.51, 127.73, 125.17, 124.99, 118.39, 51.95, 28.37. HRMS (*m/z*): calcd. for C11H9N4OCl [M+H]^+^: 249.0538; found, 249.0538.

### Compound 36

(5-methylindolin-1-yl)(1*H*-1,2,3-triazol-1-yl)methanone.

Compound **36** was synthesized as the same method as compound **17**. The carbamoyl chloride intermediate was obtained with 5-methylindoline (300 mg, 2.3 mmol), *N*,*N*-diisopropylethylamine (873 mg, 6.8 mmol), triphosgene (334 mg, 1.1 mmol) and THF (5 mL). Then the compound **35** (90 mg, 18%) was obtained as a white solid with the carbamoyl chloride intermediate, THF (5 mL), *N*,*N*-diisopropylethylamine (873 mg, 6.8 mmol), 1*H*-1,2,3-triazole (187 mg, 2.7 mmol) and DMAP (cat.). 1H NMR (500 MHz, CDCl_3_): δ 8.35 (d, *J* = 1.1 Hz, 1H), 7.99 (br s, 1H), 7.76 (d, *J* = 1.1 Hz, 1H), 7.14-7.05 (m, 2H), 4.65 (t, *J* = 8.3 Hz, 2H), 3.21 (t, *J* = 8.3 Hz, 2H), 2.36 (s, 3H). ^13^C NMR (125 MHz, CDCl_3_): δ 145.72, 139.54, 135.32, 132.84, 132.25, 128.18, 125.60, 124.90, 117.15, 51.84, 28.50, 21.04. HRMS (*m/z*): calcd. for C12H12N4O [M+Na]^+^: 251.0903; found, 251.0910.

### Compound 37

(6-nitroindolin-1-yl)(1*H*-1,2,3-triazol-1-yl)methanone.

Compound **37** was synthesized as the same method as compound **17**. The carbamoyl chloride intermediate was obtained with 6-nitroindoline (1.00 g, 6.1 mmol), *N*,*N*-diisopropylethylamine (2.36 g, 18.3 mmol), triphosgene (723 mg, 2.4 mmol) and THF (5 mL). Then the compound **37** (531 mg, 34%) was obtained as a yellow solid with the carbamoyl chloride intermediate, THF (5 mL), *N*,*N*-diisopropylethylamine (2.36 g, 18.3 mmol), 1*H*-1,2,3-triazole (505 mg, 7.3 mmol) and DMAP (cat.). 1H NMR (500 MHz, CDCl_3_): δ 8.94 (br s, 1H), 8.42 (d, *J* = 1.1 Hz, 1H), 8.06 (dd, *J* = 8.2, 2.0 Hz, 1H), 7.80 (d, *J* = 1.1 Hz, 1H), 7.42 (d, *J* = 8.2 Hz, 1H), 4.85 (t, *J* = 8.3 Hz, 2H), 3.38 (t, *J* = 8.3 Hz, 2H). ^13^C NMR (125 MHz, CDCl_3_): δ 148.03, 146.07, 143.21, 139.34, 133.17, 125.20, 125.12, 120.91, 112.74, 52.38, 28.62. HRMS (*m/z*): calcd. for C11H9N5O3 [M+H]^+^: 260.0778; found, 260.0780.

### Compound 38

(6-chloroindolin-1-yl)(1*H*-1,2,3-triazol-1-yl)methanone.

Compound **38** was synthesized as the same method as compound **17**. The carbamoyl chloride intermediate was obtained with 6-chloro-2,3-dihydro-1h-indole (376 mg, 2.5 mmol), *N*,*N-*diisopropylethylamine (949mg, 7.3mmol), triphosgene (291 mg, 1.0 mmol) and THF (5 mL). Then the compound **38** (164 mg, 27%) was obtained as a white solid with the carbamoyl chloride intermediate, THF (5 mL), *N*,*N*-diisopropylethylamine (949mg, 7.3mmol), 1*H*-1,2,3-triazole (203 mg, 2.9 mmol) and DMAP (cat.). 1H NMR (500 MHz, CDCl_3_): δ 8.37 (d, *J* = 1.3 Hz, 1H), 8.14 (br s, 1H), 7.78 (d, *J* = 1.3 Hz, 1H), 7.19 (d, *J* = 8.0 Hz, 1H), 7.13 (dd, *J* = 8.0, 1.7 Hz, 1H), 4.72 (t, *J* = 8.3 Hz, 2H), 3.23 (t, *J* = 8.3 Hz, 2H). ^13^C NMR (125 MHz, CDCl_3_): δ 145.92, 143.09, 133.34, 133.01, 130.60, 125.60, 125.46, 125.07, 117.93, 52.37, 28.14. HRMS (*m/z*): calcd. for C11H9N4OCl [M+H]^+^: 249.0538; found, 249.0541.

### Compound 39

(6-methylindolin-1-yl)(1*H*-1,2,3-triazol-1-yl)methanone.

Compound **39** was synthesized as the same method as compound **17**. The carbamoyl chloride intermediate was obtained with 6-methylindoline (149 mg, 1.1 mmol), *N*,*N*-diisopropylethylamine (434 mg, 3.4 mmol), triphosgene (133 mg, 0.4 mmol) and THF (5 mL). Then the compound **38** (164 mg, 27%) was obtained as a white solid with the carbamoyl chloride intermediate, THF (5 mL), *N*,*N*-diisopropylethylamine (434 mg, 3.4 mmol), 1*H*-1,2,3-triazole (93 mg, 1.3 mmol) and DMAP (cat.). 1H NMR (500 MHz, CDCl_3_): δ 8.35 (d *J* = 1.1 Hz, 1H), 7.94 (br s, 1H), 7.70 (d, *J* = 1.1 Hz, 1H), 7.16 (d, *J* = 7.5 Hz, 1H), 6.97 (d, *J* = 7.5 Hz, 1H), 4.65 (t, *J* = 8.5 Hz, 2H), 3.16 (t, *J* = 8.5 Hz, 2H), 2.40 (s, 3H). ^13^C NMR (125 MHz, CDCl_3_): δ 145.88, 141.99, 137.70, 132.88, 129.23, 126.21, 124.92, 124.58, 118.09, 52.13, 28.21, 21.62. HRMS (*m/z*): calcd. for C12H12N4O [M+Na]^+^: 251.0903; found, 251.0907.

### Compound 40

(7-nitroindolin-1-yl)(1*H*-1,2,3-triazol-1-yl)methanone.

Compound **40** was synthesized as the same method as compound **17**. The carbamoyl chloride intermediate was obtained with 7-nitroindoline (929 mg, 5.7 mmol), *N*,*N*-diisopropylethylamine (2.19 g, 17.0 mmol), triphosgene (672 mg, 2.3 mmol) and THF (15 mL). Then the compound **40** (51 mg, 3%) was obtained as a white solid with the carbamoyl chloride intermediate, THF (15 mL), *N*,*N*-diisopropylethylamine (2.19 g, 17.0 mmol), 1*H*-1,2,3-triazole (469 mg, 6.8 mmol) and DMAP (cat.). 1H NMR (500 MHz, CDCl_3_): δ 8.31 (d, *J* = 1.1 Hz, 1H), 7.82 (dd, *J* = 7.7, 1.1 Hz, 1H), 7.77 (d, *J* = 1.1 Hz, 1H), 7.55 (dd, *J* = 7.4, 1.1 Hz, 1H), 7.30 (t, *J* = 7.7 Hz, 1H), 4.68 (t, *J* = 8.0 Hz, 2H), 3.31 (t, *J* = 8.0 Hz, 2H). ^13^C NMR (125 MHz, CDCl_3_): δ 147.36, 140.40, 136.95, 134.62, 133.47, 129.59, 126.03, 124.79, 123.36, 54.26, 29.42. HRMS (*m/z*): calcd. for C11H9N5O3 [M+Na]^+^: 282.0598; found, 282.0602.

### Compound 41

(7-chloroindolin-1-yl)(1*H*-1,2,3-triazol-1-yl)methanone.

Compound **41** was synthesized as the same method as compound **17**. The carbamoyl chloride intermediate was obtained with 7-chloroindoline (713 mg, 4.6 mmol), *N*,*N*-diisopropylethylamine (1.80 g, 13.9 mmol), triphosgene (551 mg, 1.9 mmol) and THF (5 mL). Then the compound **40** (587 mg, 51%) was obtained as a white solid with the carbamoyl chloride intermediate, THF (5 mL), *N*,*N*-diisopropylethylamine (1.80 g, 13.9 mmol), 1*H*-1,2,3-triazole (385 mg, 5.6 mmol) and DMAP (cat.). 1H NMR (500 MHz, CDCl_3_): δ 8.34 (d, *J* = 1.1 Hz, 1H), 7.78 (d, *J* = 1.1 Hz, 1H), 7.28 (d, *J* = 7.4 Hz, 1H), 7.22 (dd, *J* = 7.4, 1.1 Hz, 1H), 7.14 (t, *J* = 7.7 Hz, 1H), 4.46 (t, *J* = 7.7 Hz, 2H), 3.20 (t, *J* = 7.7 Hz, 2H). ^13^C NMR (125 MHz, CDCl_3_): δ 146.32, 139.04, 136.70, 133.40, 129.30, 127.11, 124.48, 124.29, 123.26, 54.94, 30.50. HRMS (*m/z*): calcd. for C11H9N4OCl [M+Na]^+^: 271.0357; found, 271.0362.

### KK182 (42)

The mixture of (1*H*-1,2,3-triazol-1-yl)(4-(4-(trifluoromethyl)benzyl)piperazin-1-yl)methanone (N1-isomer) and (2*H*-1,2,3-triazol-1-yl)(4-(4-(trifluoromethyl)benzyl)piperazin-1-yl)methanone (N2-isomer).

KK182 (**42**) was synthesized as the same method as compound **17**. The carbamoyl chloride intermediate was obtained with 1-(4-trifluoromethylbenzyl)piperazine (1 g, 4.1 mmol), *N*,*N-*diisopropylethylamine (1.59 g, 12.3 mmol), triphosgene (486 mg, 1.6 mmol) and THF (5 mL). Then the compound **42** with the carbamoyl chloride intermediate, THF (5 mL), *N*,*N-*diisopropylethylamine (1.59 g, 12.3 mmol), 1*H*-1,2,3-triazole (339 mg, 4.9 mmol) and DMAP (cat.). Then the mixture was further purified on a silica gel column (Hexane:CH_2_Cl_2_:EtOAc = 1:4:0.4 to 0:4:0.4) giving KK182-N1 (260 mg, 19%) as a white solid and KK094-N2 (97 mg, 14%) as a white solid.

### KK 182-N1

^1^H -NMR (500 MHz, CDCl_3_) δ 8.18 (d, *J* = 1.3 Hz, 1H), 7.73 (d, *J* = 1.3 Hz, 1H), 7.60 (d, *J* = 8.1 Hz, 2H), 7.47 (d, *J* = 8.1 Hz, 2H), 4.10-3.71 (m, 4H), 3.62 (d, *J* = 5.3 Hz, 2H), 2.71-2.49 (m, 4H). ^13^C -NMR (126 MHz, CDCl_3_) δ 148.05, 141.61, 132.91, 129.67(q, *J* = 32.4 Hz), 129.16, 125.36, 125.32, 124.11(q, *J* = 271.8 Hz), 62.07, 52.98, 52.39, 47.86, 45.53. HRMS (*m/z*): calcd. for C_15_H_16_N_5_OF_3_ [M+H]^+^: 340.1380; found, 340.1392.

### KK 182-N2

^1^H -NMR (500 MHz, CDCl_3_) δ 7.81 (s, 2H), 7.59 (d, *J* = 8.0 Hz, 2H), 7.47 (d, *J* = 8.0 Hz, 2H), 3.95-3.64 (4H), 3.61 (s, 2H), 2.72-2.37 (m, 4H). ^13^C -NMR (126 MHz, CDCl_3_) δ 148.95, 141.77, 136.01, 129.64 (q, *J* = 32.4 Hz), 129.13, 125.32 (q, *J* = 3.8 Hz), 124.13 (q, *J* = 272.8 Hz), 62.11, 52.74, 47.67, 45.28. HRMS (*m/z*): calcd. for C_15_H_16_N_5_OF_3_ [M+H]^+^: 340.1380; found, 340.1390.

